# Evidence that interglomerular inhibition generates non-monotonic concentration-response relationships in mitral/tufted glomeruli in the mouse olfactory bulb

**DOI:** 10.1101/2025.02.28.640652

**Authors:** Lee Min Leong, David Wharton, Narayan Subramanian, Bhargav Karamched, Richard Bertram, Douglas A. Storace

## Abstract

Animals can recognize and discriminate between different odors and the same odor over a range of concentrations. Processing within the mouse olfactory bulb (OB) may be involved, yet the underlying mechanisms remain unclear. Each olfactory receptor neuron (ORN) type maps to the OB in olfactory receptor specific channels called glomeruli where they connect with the dendrites of mitral/tufted cells (MTCs), which project their axons to the rest of the brain. Differences between input and output define the functions carried out by a brain region. Using *in vivo* dual-color 2-photon Ca^2+^ imaging from the ORNs and MTCs innervating the same glomeruli in the mouse OB, we identified a novel MTC response type with a non-monotonic concentration-response relationship. We used mathematical modeling to demonstrate that non-monotonic response types are consistent with a form of interglomerular processing, which we propose is a mechanism to facilitate odor discrimination and the ability to achieve concentration-invariant odor perception.

**Graphical Abstract:** The olfactory bulb input-output transformation was imaged using dual-color 2-photon imaging. (*Top*) Cartoon and histological examples of labeling the olfactory bulb input and output using spectrally distinct optical sensors. (*Middle*) Modeling and experimental work reveal that input neurons that respond with primarily monotonic concentration-response relationships are transformed into a mix of monotonic and non-monotonic responses. (*Bottom*) Olfactory bulb output state space for a population of 3 glomeruli with exclusively monotonic responses (black), and with both monotonic and non-monotonic responses (red). Each trajectory represents a cloud of points that reflects variations in the fluctuations of molecular components present in natural odor stimuli. Monotonic responses alone cluster together, while multiple response types broaden coverage of MTC state space. Greater coverage of state space makes it easier to discriminate one odor from another, even though the dimension of olfactory bulb state space is larger than 3.

**Key Points:**

- Although the olfactory bulb is the first stage of olfactory sensory processing, its role in transforming sensory information remains poorly understood.
- Different olfactory receptor neuron types map to the olfactory bulb in olfactory receptor specific channels called glomeruli, where they interact with the dendrites of mitral/tufted cells, which project to the rest of the brain.
- We used dual-color 2-photon Ca^2+^ imaging to image the input-output transformation olfactory receptor neuron (ORN) and mitral/tufted (MTC) glomeruli across a wide concentration range.
- The ORN input to all tested glomeruli responded to increasing concentration changes with monotonic increases, while the corresponding MTCs outputs of each glomerulus responded with monotonic and non-monotonic changes.
- The results were consistent with a mathematical model of the olfactory bulb incorporating feed-forward and lateral inhibition.

## Introduction

The ability for an animal to detect and recognize odors in natural stimuli is initiated by the binding of an odor ligand to olfactory receptor neurons (ORNs), each of which express one kind of olfactory receptor (OR) protein (out of ∼1000) and have a distinct affinity for each odor (Buck & Axel, 1991; Malnic *et al*., 1999; Reisert & Matthews, 1999; Araneda *et al*., 2000; Xu *et al*., 2020; Zak *et al*., 2020). Odors evoke varying degrees of activity across the receptor population that must be related to perception. The relationship between ORs and their ligands follows classical ligand-receptor pharmacology such that activation of each OR type increases monotonically and saturates across a relatively narrow range of ligand concentration.

This has an important implication for olfactory processing and perception: the population ORN response to an odor will change when it is experienced at different concentrations or in the presence of other odors. Low concentrations of an odor will activate receptors with high sensitivity to that odor, while higher concentrations of the same odor will saturate high affinity ORs and recruit OR types with lower sensitivity to that odor (Wachowiak & Cohen, 2001; Storace & Cohen, 2017; Storace *et al*., 2019; Hu *et al*., 2020). Without further processing, an organism could not recognize or discriminate between odors in a concentration-invariant manner.

Each ORN type provides input to the olfactory bulb (OB) in OR specific channels called glomeruli (Buck & Axel, 1991; Malnic *et al*., 1999). Mitral and tufted cells (MTCs) extend their apical dendrite into a single glomerulus and project their axons to the rest of the brain (Nagayama *et al*., 2010; Igarashi *et al*., 2012; Chen *et al*., 2022). Therefore, each glomerulus contains the input-output transformation for a given OR type, which can be influenced by a complex synaptic network that includes inhibitory interneurons, and feedback from other brain regions (Cleland & Sethupathy, 2006; Parrish-Aungst *et al*., 2007; Cleland, 2010; Nagayama *et al*., 2014). Comparisons between input and output define the function(s) carried out by a brain region (Storace & Cohen, 2017, 2021).

Using *in vivo* dual-color 2-photon Ca^2+^ imaging from the ORNs and MTCs innervating the same glomeruli in response to odors presented across a wide concentration range, we identified a novel non-monotonic response type present in MTCs. In nearly half of MTC glomeruli, increases in odor concentration evoked increasing response amplitudes to a point at which they began to decrease. Because non-monotonic concentration-response relationships are unexpected in a system driven by ORN input, they indicate a transformation occurring within the OB that has implications for coding principles and the mechanisms by which they arise.

The OB network can transform ORN signals using different mechanisms that include intraglomerular processing which reflects processing within a single glomerulus, and interglomerular processing which incorporate lateral connections across glomeruli. We used mathematical modeling to demonstrate that the non-monotonic concentration-response relationships that increase then decrease reflect a form of interglomerular processing in which the MTC output reflects a combination of its afferent input, as well as the collective activity across a population (Cleland *et al*., 2007; Olsen *et al*., 2010; Carandini & Heeger, 2011).

Glomeruli within the same field of view exhibited odor-specific monotonic and non-monotonic concentration-response relationships, and higher odor concentrations broadly increased the magnitude of suppression across the OB. Comparisons between ORNs and MTCs revealed that monotonic and non-monotonic MTCs originated from ORNS with low and high sensitivity to the odor, respectively, and that suppressed MTC glomeruli are innervated by the least sensitive ORNs.

Therefore, non-monotonic MTC concentration-response relationships are widespread and the natural consequence of ORNs with a small dynamic range and interglomerular inhibition. As the concentration of a particular odor is increased, ORNs with high sensitivity to that odor will saturate, while less sensitive ORNs will become activated. Similarly, ORNs with a low sensitivity to that odor will be suppressed by rising levels of lateral inhibition from glomeruli innervated by more sensitive ORNs. The consequence is that MTCs innervated by high sensitivity ORNs will be weakened by rising levels of lateral inhibition, and those innervated by low sensitivity ORNs will be suppressed. Our results provide direct evidence of a functional transformation occurring across the OB network. We propose that the presence of non-monotonic concentration-response relationships provides a mechanism to facilitate odor discrimination and the ability to achieve concentration-invariant odor detection.

## Methods

A subset of these data were included in a previous publication that analyzed the adapting properties of MTC odor responses (Subramanian *et al*., 2025).

### Ethical approval

All experiments were carried out according to the procedures and guidelines approved by the Florida State University Animal Care and Use committee under ethics approval reference #202110074.

### Mathematical Modeling

The model is based on ideas described in (Cleland & Sethupathy, 2006) and incorporates two forms of inhibition: local and lateral. The output of the ORNs is dependent upon the odorant concentration, [odorant], and is described mathematically by the sigmoidal Hill function:

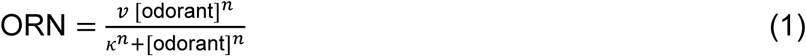

All variables and parameters are dimensionless, and [odorant] ranges from 0 to 1. The parameter 𝜐 sets the maximum value of ORN and is assumed to be the same for all ORNs. The other two parameters are the half-activation constant, 𝜅, and the Hill coefficient, *n*, which sets the steepness of the ORN’s response (higher values of *n* yield steeper response functions). To generate a large (*N* = 949) heterogeneous population of ORNs, *N* combinations of parameter values were sampled from a uniform distribution on the intervals of (0,2) for 𝜅 and (1,4) for *n*. The ORN signal is then subject to presynaptic inhibition from other glomeruli before entering the target glomerulus, producing a “normalized ORN” value 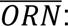

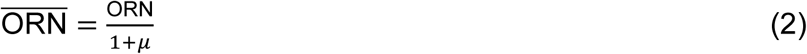

where 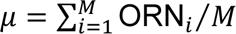 is the average ORN activity over a population ORNs. We consider two cases. In the first, each ORN is normalized by a randomly selected population of *M* = 50 ORNs. In the second case, each ORN is normalized by all other *M* = 948 ORNs. This normalized output of the ORN is then the input to periglomerular and MTCs, reflected in the variables PG and MTI, respectively. The circuitry involved in this lateral inhibition is not specified in the model, and could reflect the action of short axon cells onto external tufted cells, which then serve as input to MTCs and PG cells, as previously described (McGann, 2013; Banerjee *et al*., 2015). The PG and MTI variables are described with increasing Hill functions:

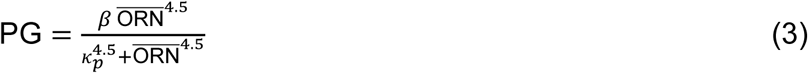

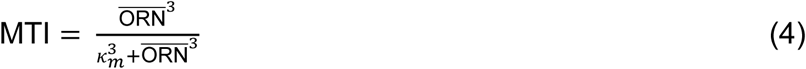

where *β*, 𝜅_*p*_, and 𝜅_*m*_ are parameters that are assumed to be the same for each PG and MTI. They are set so that PG begins to activate at lower values of 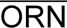 than MTI and saturates at a lower value (saturation value of *β*) than MTI (saturation value of 1). The higher affinity of PG cells is a key element of the half-hat coding hypothesized by Cleland (Cleland & Sethupathy, 2006; Cleland, 2010). The output of each glomerulus, MTC output, is then the difference between MTI and the inhibition from PG cells,

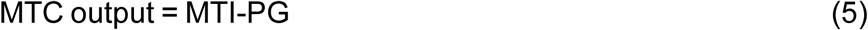

Parameter values are given in **Table 1**.

**Table 1:** Parameter values used in model simulations.

| Symbol | Value(s) | Description |
| --- | --- | --- |
| [odorant] | [0,1] | concentration of input stimuli |
| $v$ | 2 | Maximum ORN value |
| $n$ | random(1,4) | ORN Hill coefficient |
| $\kappa$ | random(0,2) | ORN half-activation value |
| $\beta$ | 0.6 | Maximum PG value |
| $\kappa_p$ | 0.5 | PG half-activation value |
| $\kappa_m$ | 1 | MTI half-activation value |

### Transgenic mice

GCaMP6f was targeted to MTC glomeruli by mating the Ai148 GCaMP6f transgenic reporter line (Jax stock #030328) to the Tbx21-cre transgenic line (Jax stock #024507) (Mitsui *et al*., 2011; Daigle *et al*., 2018; Subramanian *et al*., 2025). GCaMP6s was targeted to olfactory receptor neuron (ORN) glomeruli by mating the tetO-GCaMP6s transgenic reporter line (Jax stock #024742) to the OMP-tTA transgenic line (Jax stock #017754) (Huang *et al*., 2022; Subramanian *et al*., 2025). This OMP-tTA-tetO-GCaMP6s line was mated with the Tbx21-cre transgenic line (Jax stock #024507) to create a triple transgenic line (OMP-tTA-tetO-GCaMP6s Tbx21-cre) expressing GCaMP6s in the ORN glomeruli and allowing for cre-dependent expression in the MTC glomeruli. GCaMP8m was targeted to MTC glomeruli by mating the TIGRE2-jGCaMP8m-IRES-tTA2-WPRE transgenic reporter line (Jax stock # 037718) to the Tbx21-cre transgenic line (Jax stock #024507) (Mitsui *et al*., 2011; Daigle *et al*., 2018; Subramanian *et al*., 2025). Genotyping was performed by Transnetyx (Cordova, TN) and male and female adult offspring that expressed eGFP and Cre recombinase for MTC glomeruli or eGFP, tTA and Cre recombinase for ORN glomeruli were used for experiments. We further confirmed that the fluorescent proteins were expressed in the excepted cell population in histological samples (described under Histology).

### Surgical procedures

All procedures were approved by the Florida State University Animal Care and Use Committee. All mice included in this study were maintained in the FSU animal vivarium on a 12h/12h light/dark rhythm with ad libitum access to food and water. Male and female adult (> 21 days) transgenic mice were anesthetized using ketamine/xylazine (90 / 10 mg/kg, IP, Zoetis, Kalamazoo, MI), placed on a heating pad and had ophthalmic ointment applied to their eyes. Mice were given a pre-operative dose of carprofen (20 mg/kg, SC, Zoetis, Kalamazoo, MI), atropine (0.2 mg/kg, IP, Covetrus, Dublin, OH), dexamethasone (4 mg/kg, IP, Bimeda, La Sueur, MN), and bupivacaine (1.5 mg/kg, SC, Hospira, Lake Forest, IL). Fur was removed from the top of the skull using a depilatory agent and rinsed, after which the skin was scrubbed with 70% isopropyl alcohol and iodine (Covidien, Mansfield, MA). An incision was made to remove the skin over the skull and blunt dissection was used to remove the underlying membrane. Dental cement (Metabond, Covetrus, Dublin, OH) was used to attach a custom headpost to the skull, which was held using a custom headpost holder. After the cement finished drying, the bone above the OB was either thinned using a dental drill (Osada, XL-230, Los Angeles, CA) and covered with cyanoacrylate to improve optical clarity or was removed and replaced with cover glass.

For the dual color input-output experiments, stereotaxic injections were carried out immediately after the craniotomy. 500 nl of AAV1. Syn.Flex.NES-jRCaMP1b.WPRE.SV40 (Addgene, 100850-AAV1, Watertown, MA) was injected into the right olfactory bulb of OMP-tTA-tetO-GCaMP6s Tbx21-cre triple transgenic mice before sealing the cranial window with a #1 cover glass. Upon completion of the surgery, the mouse was allowed to recover on a heating pad until they were fully ambulatory. Animals were given a post-operative dose of carprofen (20 mg/kg, SC) at the end of the day of surgery and for at least 3 days post-operatively. The AAV was allowed to express for at least two weeks before beginning imaging experiments.

### Histology

We validated the targeting of the optical sensors used in this study based on endogenous fluorescence expression of GCaMP6f (MTCs), GCaMP6s (ORNs), GCaMP8m (MTCs) and jRCaMP1b (MTCs). Histological samples were observed using appropriate filter sets on a Zeiss Axioskop epifluorescence microscope and were imaged on a Nikon CSU-W1 spinning disk confocal microscope using a 10x 0.45 N.A., 40x 0.95 N.A. or 40x 1.15 N.A. objective lenses. Euthanasia was carried out via procedures approved by the FSU ACUC. Mice were euthanized via an IP injection of euthasol and either underwent cardiac perfusion with phosphate buffered saline and 4% paraformaldehyde or had their brains directly extracted and post-fixed in 4% paraformaldehyde before being cut on a vibratome in 40 µm sections (Leica VT1000S, Deer Park, IL). OB sections were mounted on slides and were coverslipped using Fluoromount-G containing DAPI (SouthernBiotech, Birmingham, AL).

### 2-photon imaging

Imaging was performed using a Sutter MOM 2-photon microscope equipped with an 8 kHz (30.9 Hz) resonant scanner (Cambridge Technology, USA) and an emission pathway equipped with a GaAsP PMT (#H10770PA-40-04, Hamamatsu, Japan). Laser excitation was provided using either an Alcor 920-2W laser (920 nm for GCaMP imaging) with power modulated by an internal acousto-optic modulator or a Spectra-Physics DS+ between 940-980 nm for GCaMP imaging or 1100 nm for jRCaMP1b imaging, with power modulated by a Pockels cell (Model #350-105-02, Conoptics, Danbury, CT). Imaging was performed using a Nikon 16x 0.8 N.A. or a 10x 0.5 N.A. objective lens (Yu *et al*., 2024). Laser power was confirmed to be less than 150 mW at the output of the objective lens measured using a power meter (Newport 843-R) for scanning areas ranging between 711 µm^2^ - 1138 µm^2^.

### Odorant delivery

Methyl valerate (99% pure, CAS #624-24-8, Sigma-Aldrich, #148997), isoamyl acetate (99% pure, CAS #123-92-2, Thermo Scientific #150662500), benzaldehyde (> 99.5% purity, CAS #100-52-7, Sigma Aldrich #418099), acetophenone (99% pure, CAS #100-52-7, Fisher Scientific #AAA12727AP), and 2-phenylethanol (> 99% CAS #60-12-8, Sigma-Aldrich #77861) were used at concentrations between 0.05 and 5.5 % of saturated vapor. The olfactometer design involved air being pushed through vials of pure odor using a syringe pump (NE-1000, PumpSystems, Farmingdale, NY) running at different flow rates (0.25 – 28 ml /min) (Subramanian *et al*., 2025). This odor stream underwent an initial air dilution with a lower flow rate of clean air (30 ml/min). The resulting odorized air stream connected to a dual 3-way solenoid valve (360T041, NResearch, West Caldwell, NJ) which was connected to an exhaust, a clean air stream and a Teflon delivery manifold which served as the final delivery apparatus placed in front of the mouse’s nose. The delivery manifold had a higher flow rate of clean air constantly flowing through it (450 ml/min), which provided a second air dilution. At rest, the solenoid sent the odorized air stream to the exhaust, and the clean air stream to the animal; odor delivery occurred by triggering the solenoid which caused the odor to be injected into the delivery manifold.

For the dual color input-output experiments, the odorants methyl valerate, isoamyl acetate, acetophenone, and methyl salicylate (99% pure, CAS#119-36-8, Sigma-Aldrich #M2047) were diluted in mineral oil to 0.002%, 0.008%, 0.046%, 0.278%, 1.667% and 10%. The olfactometer setup uses an air pump that is constantly delivering clean air to two mass flow controllers (MC-100SCCM, and MC-1SLPM, Alicat, Tucson, AZ).

The odor mass flow controller (MC-100SCCM) controls the air flow that passes through the odor vials (50 ml/min for this study), the dilution mass flow controller (MC-1SLPM) provides a higher flow rate of air (450 ml/min for this study) that dilutes the odorized air (i.e., the odors were delivered at 10% saturated vapor). The mass flow controllers are each connected to a 4-valve manifold (360T081, NResearch, West Caldwell, NJ) where the odor vials are connected in between. At any given time, only one of the four pairs of manifold valves open to allow for the odor air stream to blend with the dilution air stream to deliver a particular concentration of the mineral oil diluted odors. The resulting odorized air stream connected to a dual 3-way solenoid valve (360T041, NResearch, West Caldwell, NJ) which was connected to an exhaust, a clean air stream and a Teflon delivery manifold which served as the final delivery apparatus placed in front of the mouse’s nose. The delivery manifold has a separate air pump that delivers clean air at a flow rate that is matched to the flow rate output of the mass flow controllers. The 3-way solenoid valve sends the odorized air stream to the exhaust and the clean air stream to the mouse prior to odor trigger. Triggering the 3-way solenoid causes the odor to be injected into the delivery manifold. The odor delivery time-course for both olfactometer setups were confirmed using a photoionization detector (200C, Aurora Scientific, Aurora, ON) (Storace & Cohen, 2017, 2021; Subramanian *et al*., 2025).

### Imaging procedures

Prior to data collection mice were positioned underneath the microscope and the headpost holder angle was adjusted to optimize the imaging field of view. During data collection, awake head-fixed mice were placed underneath the microscope objective with the olfactometer and either a thermocouple (Omega 5TC-TT-K-36-36, Newark) near its nose, or a pressure sensor embedded in the olfactometer to measure respiration. The signals from the respiration sensor were amplified and low-pass filtered using a differential amplifier (Model 3000, AM-Systems, Sequim, WA), which was recorded by the imaging system. For GCaMP6f imaging experiments (**Figures 2-6**), odors were delivered at concentrations between 0.05 – 5.5 % of saturated vapor in trials separated by a minimum of 3 minutes. For glomerular imaging using jGCaMP8m (MTCs), jRCaMP1b (MTCs) and GCaMP6s (ORNs), odors were delivered in 6 concentration steps between 0.002% - 10% of pure odor diluted in mineral oil at 10% of saturated vapor, and air trials in which the odor stream was passed through a previously unused vial. For each animal, we prioritized measuring multiple repetitions for each concentration for a particular odor before a second odor was attempted.

### Data analysis

#### Frame Subtraction Analysis

The mean fluorescence and frame subtraction images are from the average of at least two single trials (**Figure 2A, 7C**). The mean fluorescence images are generated from the average of all the frames during the imaging trial, or all the frames prior to odor stimulation. The frame subtraction images were generated by subtracting an average of the 19 frames during odor stimulation from the average of 9 frames prior to the stimulus. The resulting image underwent two passes of a low-pass spatial filter, and the fluorescence (F) values were converted to ΔF/F by dividing the fluorescence value of each pixel by the mean of at least 60 consecutive frames in the image stack in Turbo-SM. The intensity scale range is fixed to the same minimum and maximum range for all concentrations for each odor (**Figure 2**).

#### Processing and segmentation

Following data acquisition, the raw image files were spatially and temporally averaged from 512x512 pixels sampled at 30.9 Hz to 256x256 pixels sampled at 7.72 Hz. The resulting data were exported to TIFF format for all subsequent analysis. Occasional recordings with motion artifact that made it impossible to interpret the measurements were discarded from subsequent analysis. Glomerular regions of interest were manually segmented in custom software (Turbo-SM, SciMeasure, Decatur, GA) and were identified based on their morphological properties in the mean fluorescence, and functional responses in a frame subtraction. The pixel areas containing the regions of interest were saved and the fluorescence time course values from each region of interest were extracted for subsequent analysis. Fluorescence time course values were converted to ΔF/F by dividing each trace by the mean of the frames prior to odor stimulation. Odor response amplitudes and corresponding Z-Scores were calculated as the largest difference between a 1200 msec window during the odor presentation and the time prior to odor stimulation.

#### Other analyses

Population descriptive statistics were quantified in responsive glomeruli which were defined as those in which an odor evoked a minimum of a 5 standard deviation change from baseline at any concentration (**Figure 2F**). Threshold and best response are defined as the lowest concentration evoking a response, and the concentration evoking the largest response, respectively (**Figure 2G-H**).

For all other analyses, glomeruli were included that responded with a minimum of 3 standard deviations above baseline at some concentration. The response category assigned to each MTC glomerulus was defined based on a combination of visual inspection of the mean concentration-response curve, the corresponding single trials, and the Z-score change in response to the odor stimulus. Excited and suppressed glomeruli were defined based on having clear odor-locked increases or decreases that could be clearly identified in the mean and single trial fluorescence time course. Population measurements of the different response categories were generated by averaging the fluorescence time course from all the glomeruli in a field of view assigned to that category (**Figure 3E-F**).

For the spatial maps of glomerular ΔF/F and categorical responses, glomeruli that did not respond at any concentration are indicated using cross-hatching (**Figure 4A-B**). For responsive glomeruli, concentrations that evoked a change of less than 2 standard deviations were assigned a value of zero to facilitate visual analysis (**Figure 4A**, *white polygons*). For the network category analysis (**Figure 4B-C**), the response category of each glomerulus was included for preparations that were tested to more than one odor. The network graph in **Figure 4C** was generated by plotting the categories of each glomerulus for different odors using the graph function in MATLAB.

The relationship between excitation and suppression was quantified by averaging the response from all glomeruli that responded to the highest concentration with excitation and suppression, respectively (**Figure 5**). The mean excited and suppressed response for each preparation are plotted together, with the responses from individual preparations connected with a line (**Figure 5E**). The correlation between these values were quantified for individual preparations and across the population using the corrcoef function in MATLAB (**Figure 5D-E**). The variance explained was quantified by binning the mean excitation and suppression value measured for each concentration (**Figure 5E**, black solid line), and fitting them with a sigmoid (**Figure 5E**, black dashed line) (Economo *et al*., 2016).

To avoid biasing the proportion of the dataset identified as non-monotonic, we analyzed the GCaMP6f dataset using two approaches. The first approach used the Monotonicity Index (MI), a metric that computes the degree of non-monotonicity in a concentration-response function (**Figure 6**) (Escabí *et al*., 2007; Higgins *et al*., 2008). The MI of individual glomeruli was calculated by computing the d-prime value by measuring the concentration-response function (CRF) in ΔF/F.

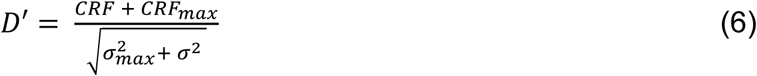

The CRF_max_ is the response at the concentration with the largest ΔF/F change from the baseline and CRF is the ΔF/F change at each concentration. σ^2^_max_ is the response variance at the highest odor concentration, while σ^2^ is the response variance at each odor concentration. MI was then computed as the minimum D’ measured at concentrations beyond the one evoking the maximum ΔF/F response. Consequently, a monotonically increasing glomerulus is assigned a value of 0, while glomeruli with non-monotonic concentration response functions are assigned negative values. Glomeruli with moderate non-monotonic shapes are closer to 0, while those with a larger decrease are assigned increasingly negative values. For the analysis in **Figure 6**, MI values were quantified for all glomeruli in the GCaMP6f data set except for exclusively suppressed glomeruli because they are assigned large MI values. The second approach quantified the general slope of the MTC concentration-response function (Shen *et al*., 2025a). The linear slope of MTC glomeruli was quantified by first performing a linear interpolation between each consecutive pair of points in the concentration-response relationship. The polyfit function in MATLAB was then used to find the average slope of each line.

half-maximum values were quantified by performing a linear interpolation of the concentration-response relationship for each glomerulus (linearinterp fittype in MATLAB) (**Figures 6-7**, vertical lines). The half-maximum value was defined as the concentration evoking the half-maximum response in the linear interpolation except for the caveat in which glomeruli with a response to the lowest concentration that was greater than the half-maximum were assigned a half-maximum value of 0.05% (e.g., **Figure 6B**, roi9).

Hill equation fits of monotonically increasing glomeruli were performed in glomeruli exhibiting MI values greater than -0.5, Hill fits were calculated using the following equation where *X* is the response (in ΔF/F) of the glomerulus at each concentration, and *k* is the half-saturating value estimated from a linear interpolation of the concentration-response function.

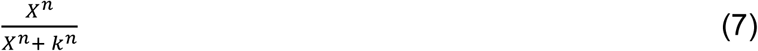

Fits were only analyzed if they met the following criteria: the Hill coefficient was between 0.75 and 6; the r^2^ value was greater than 0.9, the root mean sum of squares was less than 0.1 and the sum of square error was less than 0.1 (Zak *et al*., 2020).

The trajectory plots were generated by plotting the odor response amplitudes of 3 glomeruli exhibiting exclusively increasing responses, or a combination of all response categories using the plot3 function in MATLAB (**Figure 10**). Data are presented as mean ± standard error of the mean. All error bars represent standard error of the mean.

## Results

### A model that combines intra- and interglomerular processing predicts four types of concentration-response relationships in MTC glomeruli

We generated a model to better understand how intra- and interglomerular processing transforms the concentration-response properties of different ORN inputs (see Methods for model equations and parameter values). The output of each glomerulus (**Figure 1A**, d) was a function of its ORN input (**Figure 1A**, a), input from local interneurons that are activated by ORN input and provide inhibition onto the MTC output (**Figure 1A**, b), and inhibitory interglomerular connectivity acting presynaptically to the MTCs (**Figure 1A**, c). ORNs modeled using the Hill equation served as input both to MTCs and inhibitory intraglomerular interneurons whose activation was described by a Hill function left-shifted relative to the MTC response function (**Figure 1B**, PG). The difference between these two response functions was then taken as the output from the MTC (**Figure 1B**, MTC output). The MTC output, which is relative to the basal level of activity, has a half-hat shape that shows a dip to negative values for low ORN activation and an increase for higher ORN activation, as previously described by Cleland (**Figure 1B**, MTC output) (Cleland & Sethupathy, 2006).

**Figure 1:**
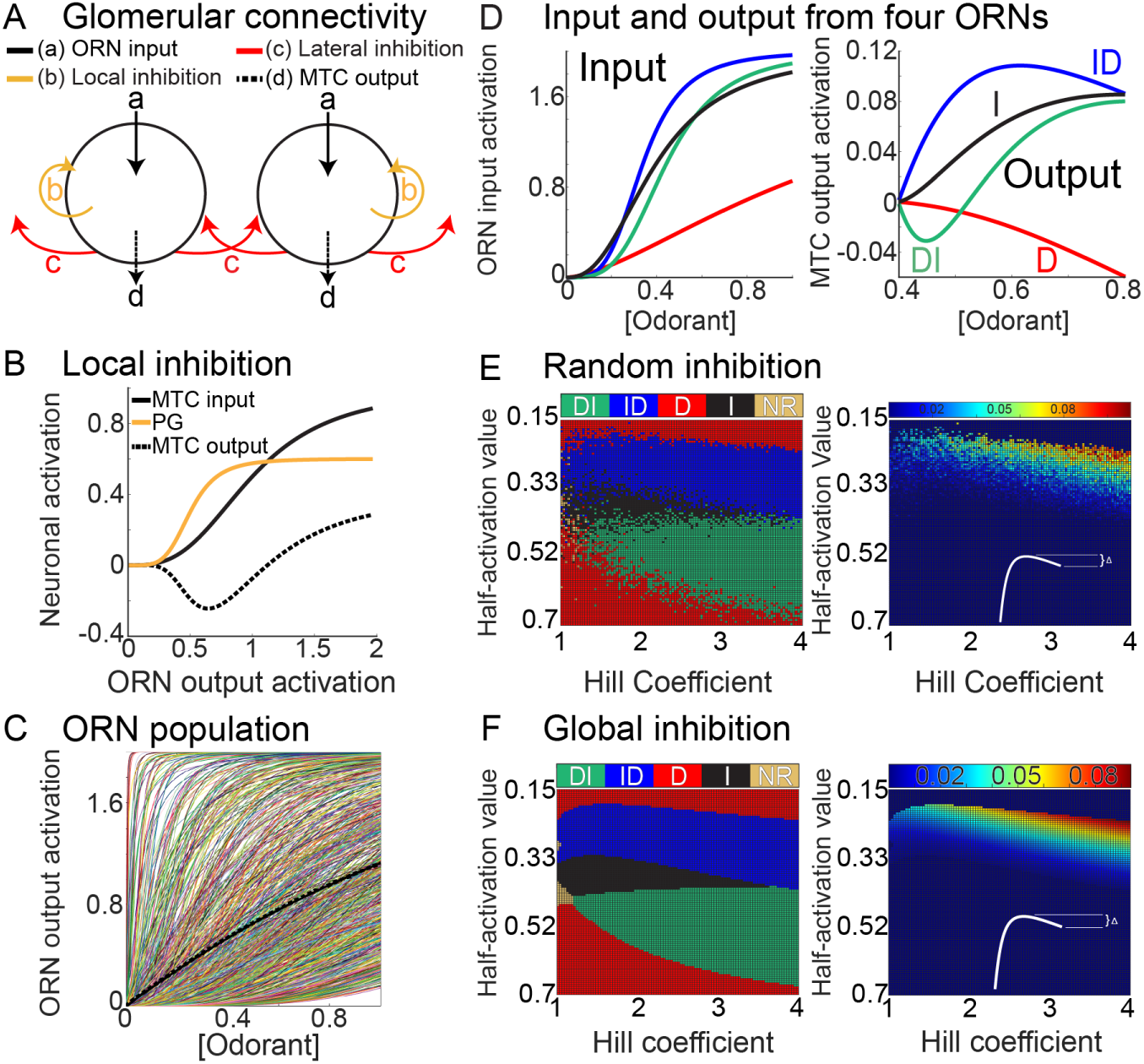
(**A**) Model of the olfactory bulb input-output transformation. (**B**) Local inhibition alone generates a half-hat response in MTCs. (**C**) The population of 949 ORNs used in the model. One example of a normalizing curve is shown (dashed). (**D**) The concentration-response relationships of four ORNs (left) and their corresponding MTC output (right) reflecting both local and lateral inhibition. Output responses were categorized based on the shape of the concentration-response relationship: I, increasing; ID, increasing then decreasing; DI, decreasing then increasing; D, decreasing. (**E**) (left) MTC responses for different ORN Hill coefficients and half-activation values, in which normalization of input into each glomerulus is through a randomly-selected subpopulation of 50 glomeruli. (right) The decrease from the maximum to the terminal odorant concentration value for combinations of Hill coefficients and half-activation values that result in an ID response. (**F**) Similar to the previous panel, but now normalization of input into each glomerulus is through all other glomeruli.

Interglomerular processing was modeled by generating a population of 949 ORNs with Hill coefficients sampled from a uniform distribution from 1 to 4, consistent with previously published work (**Figure 1C**, colored thin lines)(Firestein & Zufall, 1993; Wachowiak & Cohen, 2001; Storace & Cohen, 2017; Zak *et al*., 2020; Platisa *et al*., 2022). Odor sensitivities (their half-maximum values) were sampled from a uniform distribution on the interval between 0 and 2 (**Figure 1C**). The model is dimensionless, as are the half-maximum values for the receptors. The ORN input to the MTCs and inhibitory PG cells was divided by the mean ORN activity over a population of 50 randomly-selected ORNs to reflect lateral inhibition from a small subset of the glomeruli. This structure results in a normalization procedure in which the outputs of each MTC glomerulus are influenced by the input from a randomly-selected subset of ORN glomeruli (Carandini & Heeger, 2011). One example of a normalizing function is shown as a dashed curve in Figure 1C.

Four examples of ORN input and corresponding MTC output across a concentration range are illustrated in **Figure 1D** (the colors are matched across the two subpanels). The model output in response to four input functions could be broadly described as having concentration-response relationships that monotonically increased (I), monotonically decreased (D), transitioned from suppressed to excited (DI) or increased to a point at which further concentration increases evoked progressively smaller responses (ID) (**Figure 1D**, right subpanel).

The types of MTC responses are shown with color coding in **Figure 1E** (left) over a grid of values of the ORN Hill coefficient and half-activation constants. The ORNs with the highest sensitivities to an odor had a purely decreasing response to increasing odor concentrations (**Figure 1E**, D, top red regions). In these cases, the MTC output reached its maximum at low odor concentrations and could only decline due to lateral inhibition as other glomeruli became activated at higher concentrations. For ORNs with lower, but still relatively high sensitivities, the typical response was an initial increase followed by a decrease (**Figure 1E**, ID, blue region). In these cases, strongly activated ORNs near saturation were dominated by increasing lateral inhibition. For lower-sensitivity ORNs the response was purely increasing, particularly when the ORN response function was relatively linear (**Figure 1E**, I, black region). At even lower sensitivities the typical response was a decrease, followed by an increase (**Figure 1E**, DI, green region). In this case, inhibition dominated the MTC output prior to its strong activation at higher odor concentrations, producing a DI response. Finally, at the lowest sensitivities the response was purely decreasing, reflecting dominance of inhibition over all odor concentrations considered (**Figure 1E**, D, red region).

In principle, D responses can be generated by feed-forward suppression within a range of concentrations (e.g., **Figure 1B**, dashed line between 0 and 0.6), or in the case of spontaneously active ORNs suppressed by interglomerular inhibition. ID responses cannot, however, result from the local inhibition motif used in our model. Instead, they depend upon lateral inhibition, which we hypothesize results from higher levels of interglomerular inhibition resulting from activation of ORN glomeruli with lower sensitivities to the odor. We tested this hypothesis by excluding lateral inhibition from the model, which resulted in a loss of the ID response types (**Supplementary Figure A1A-B**).

A consequence of this framework is that the magnitude of non-monotonicity should be heterogeneous and originate from higher affinity glomerular input that saturates at lower concentrations. A quantification of the decrease in MTC activation from the maximum over the grid of ORN parameters evoking ID responses revealed a gradient of non-monotonic drops (**Figure 1E**, right). Most of the parameter values in the grid that produced a large drop in MTC activation following a peak had small ORN half-activation values indicating high sensitivity for the odor.

The structure of the interglomerular inhibitory network is currently unknown (Fantana *et al*., 2008; Banerjee *et al*., 2015; Economo *et al*., 2016). The results shown in **Figure 1E** were obtained assuming that each glomerulus is subject to lateral inhibition from a subpopulation of 50 randomly-selected glomeruli. When instead the subpopulation was increased to include all glomeruli, so that there is all-to-all coupling among the glomeruli, the results are qualitatively similar (**Figure 1F**). With this larger normalizing population, the boundaries between the different response categories become sharper, but the basic structure of the response pattern is the same. In general, we found that the main effect of reducing the size of the randomly-selected normalizing set of glomeruli was to increase the fuzziness of the boundaries between the different response regions. We also considered the case in which input from 5 and 10 randomly chosen glomeruli provided the interglomerular inhibition, which resulted in different average ORN activation curves (Eq. 2) (**Supplementary Figure A1C**). Importantly, all four types of response patterns were produced when interglomerular processing was implemented with the random subsets of ORNs (**Supplementary Figure A1D**). It is clear, then, that the four different response types are generic, while the shapes and extent of the regions in the 2-parameter grid that produce the different types of responses depend on the structure of the interglomerular inhibitory network.

### In vivo measurements of concentration-response relationships in MTC glomeruli

We tested these predictions in a data set in which odor responses were measured from MTC glomeruli across a ∼100-fold change in concentration using *in vivo* 2-photon Ca^2+^ imaging in awake mice. Imaging was performed in transgenic mice in which GCaMP6f was selectively expressed in MTCs and their corresponding apical dendrites innervating the glomerular layer (**Figure 2A**, *mean fluorescence*). We histologically confirmed targeting of GCaMP6f to MTCs in 3 preparations (*not shown, but see* (Subramanian *et al*., 2025)). Frame subtraction analyses from different preparations illustrate that odors evoked different patterns of excited and suppressed responses within the same field of view (**Figure 2A**, *white versus black regions in frame subtractions*). The specific activation pattern across the glomerular population was a complex function of odor concentration (**Figure 2A***, compare over concentration range 0.05 to 5.5%*). Many glomeruli became increasingly excited (monotonically increasing, I) or suppressed (monotonically decreasing, D) at higher concentrations (**Figure 2A**, I vs D). Some glomeruli exhibited nonlinear changes in response to increasing concentration in which they first decreased then increased (DI), while others increased and then decreased (ID) (**Figure 2A**, DI vs ID).

**Figure 2:**
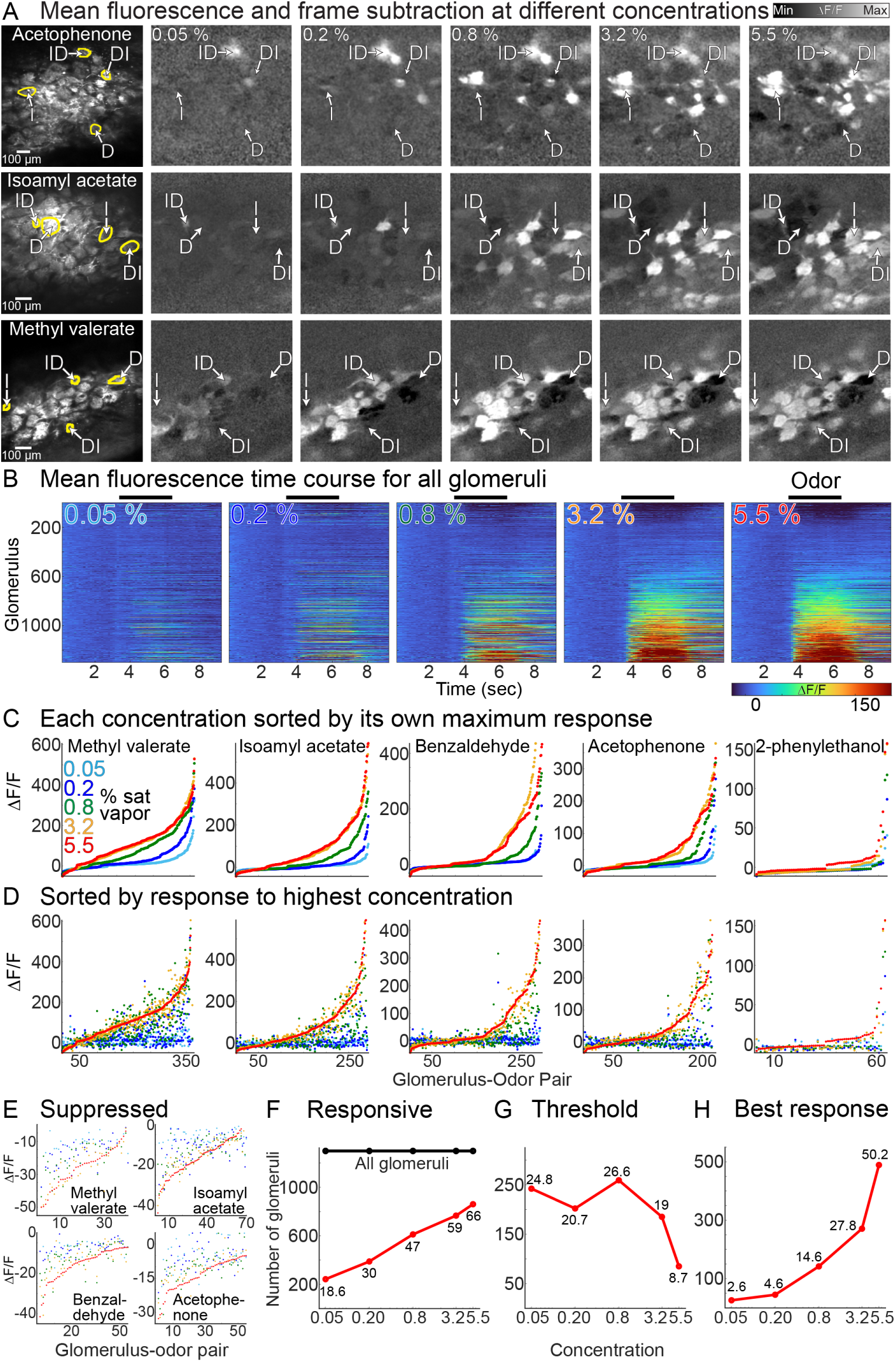
(**A**) Mean fluorescence (left subpanel) and frame subtractions illustrating MTC activation at different concentrations in 3 preparations (rows). The intensity scale is fixed within each preparation. The arrows highlight glomeruli with increasing (I), decreasing (D), increasing-decreasing (ID) and decreasing-increasing (DI) concentration-response relationships. (**B**) Fluorescence time course from all glomerulus-odor pairings. The color scale is fixed across the 5 concentrations. (**C**) Response to all glomeruli for each odor sorted by the maximum response to each concentration (colors). (**D**) Same arrangement as panel **C** except that glomeruli are sorted by the maximum response to the highest concentration. (**E**) Data from panel **D** cropped to visualize suppressed glomeruli. (**F**-**H**) Descriptive statistics. The numbers on each plot indicate the fraction of total glomeruli.

The response from each glomerulus was segmented, and the fluorescence time course and odor response amplitudes (ΔF/F) and corresponding Z-scores were calculated as the largest difference in a 1200 msec moving window between the odor stimulation period and the baseline fluorescence. We quantified the concentration-response relationships in a population of glomerulus-odor pairings that included 736 individual glomeruli in 20 different preparations (36.8 ± 3.3 glomeruli per preparation, ranging from 13-64). At least one odor was delivered to each preparation at 5 different concentrations, which included the odors methyl valerate (n = 14), isoamyl acetate (n = 7), benzaldehyde (n = 7), acetophenone (n = 5), and 2-phenylethanol (n = 1). The data set includes 6495 glomerulus-odor pairings across all tested odors and concentrations (**Figure 2B**). The fluorescence time course versus amplitude of all glomerulus-odor pairings illustrates that higher concentrations typically evoked stronger degrees of excitation and suppression in more glomeruli (**Figure 2B**), though there are instances where this is not true, as discussed next.

We examined the relationship between concentration and response magnitude at the population level by plotting the ΔF/F evoked for each glomerulus-odor pairing for different odors (**Figure 2C-D**). Sorting glomerular responses from smallest to largest for individual concentrations illustrates that relatively few glomeruli are activated at low concentrations, while the number of activated glomeruli and their overall response amplitude increased at higher concentrations (**Figure 2C**). Sorting glomeruli by their response to the highest concentration more clearly visualizes concentration-response heterogeneity across the population (**Figure 2D**). For many glomeruli, the response to the highest concentration evoked the largest amplitude response (**Figure 2D**, the red point is above the others), although other glomeruli responded with a sub-maximal response to the highest concentration (**Figure 2D**, points of other colors are above red). For glomeruli with suppressed responses, the highest concentration tended to evoke the strongest magnitude of suppression, but not always (**Figure 2E**, data are cropped from panel **D**).

We quantified these observations using a statistical threshold in which significantly responsive glomeruli were defined as responding to the odor with a 5 standard deviation change from the pre-odor frames. The number of significantly responsive glomeruli increased as a function of odor concentration (**Figure 2F**). A similar analysis was used to define the lowest concentration that evoked a significant response for each glomerulus (the “threshold” concentration), and the concentration that evoked the strongest response (the “best” concentration). Of the responsive glomeruli, approximately half had a threshold response of 0.8% of saturated vapor, while the remaining first responded to a lower or higher concentration (**Figure 2G**). About half of the glomeruli responded most strongly to the highest tested concentration, while the remainder responded most strongly to a lower concentration (**Figure 2H**). Increasing or decreasing the statistical threshold altered the number of “responsive” glomeruli in the data set but did not substantially change the mean threshold and best response of the glomeruli (*not shown*).

### Examples of different MTC response types

Representative examples from four different preparation-odor pairings illustrate that monotonic and non-monotonic glomerular concentration-response relationships were consistent across single trials and often present in the same field of view (**Figure 3A-D**, all four glomeruli in each subpanel were simultaneously imaged in the same field of view).

**Figure 3:**
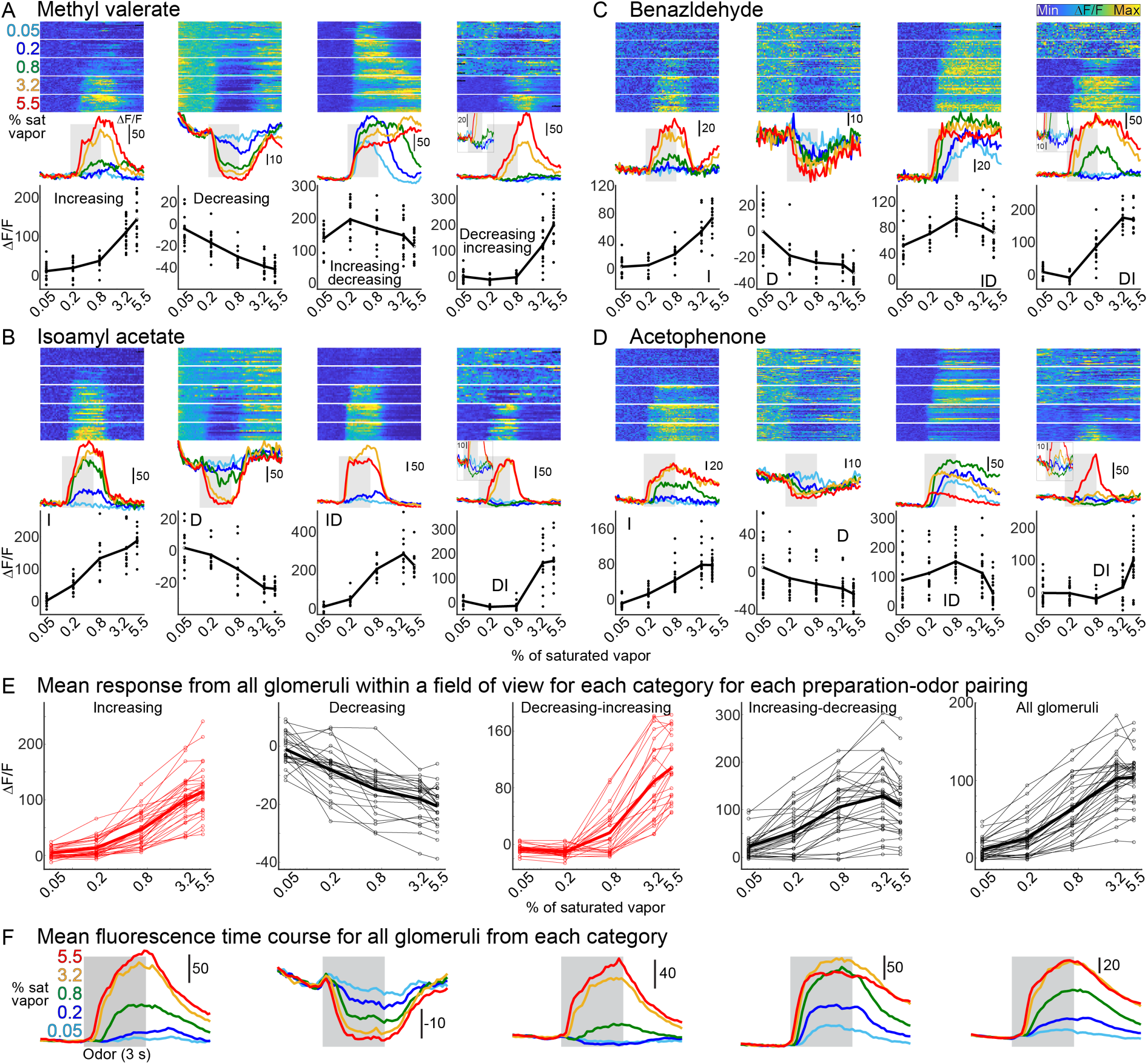
(**A**-**D**) Single trials (top), mean fluorescence time course (middle) and concentration-response relationships with points indicating the response amplitudes from the single trials for four different preparations. (**E**) Mean concentration-response relationships for each preparation-odor pairing for each response type (1st four subpanels), and across all glomeruli in each field of view (right-most subpanel). (**F**) Mean fluorescence time course from all glomeruli in each category.

Glomeruli categorized as I were present in 336 glomeruli and were defined as those that exhibited monotonically increasing response amplitudes from the lowest to the highest concentration, and which were never suppressed by the odor (**Figure 3A-D**, 1^st^ subpanel). I glomeruli could be well fit with the Hill equation with a mean Hill coefficient of 2.7 ± 0.08 (range of 0.8-5.9, included 190 glomeruli in 16 preparation-odor pairings, the remainder were excluded due to poor fits, see Methods). We observed changes consistent with D in 125 glomeruli, which were defined as those that never increased in fluorescence in response to any odor-concentration pairing, but which exhibited a clear odor-locked decrease at the highest concentration. The mean ΔF/F and Z-score of D glomeruli at the highest concentration was -21.6 ± 0.9 and 6.8 ± 0.4, respectively) (**Figure 3A-D**, 2^nd^ subpanel). Glomeruli consistent with the ID response type were also present where higher concentrations drove increased response amplitudes up to a point at which further increases caused progressively smaller, but still excited responses (**Figure 3A-D**, 3^rd^ subpanel). ID concentration-response relationships were measured in 362 glomeruli, and had a mean decrease in ΔF/F from the largest response amplitude to the response evoked at the highest concentration of 29.2 ± 1.7. Response types characteristic of DI were present in 92 glomeruli and were defined as having a suppressed response to at least one sub-maximum concentration, and an excitatory response to the highest concentration (**Figure 3A-D**, 4^th^ subpanel). For DI glomeruli, the mean ± s.e.m. ΔF/F and Z-scores at the concentration with the largest magnitude suppressed response was -13.7 ± 0.97 and 4.15 ± 0.23, respectively and occurred most commonly at 0.2% (42/92) or 0.8% (35/92) of saturated vapor. Finally, 34 glomeruli exhibited concentration-response relationships that included suppression but could not be clearly fit into any category (e.g., glomeruli that transitioned from strongly excited to suppressed, *not shown*). Thus, of the 915 glomeruli that could be categorized, 37% were of type I, 14% were of type D, 10% were of type DI, and 39% were of type ID.

Plots in which the concentration-response relationship of all glomeruli from each category were averaged together in each preparation-odor pairing illustrate that each response type was consistently present in nearly all preparations (**Figure 3E**, left 4 subpanels, each line represents the average from one preparation). Averaging all responsive glomeruli together within the same field of view demonstrates that higher concentrations drives overall increasing levels of excitation across the glomerular population (**Figure 3E**, All glomeruli, right-most subpanel), although this is often not true at the level of a single glomerulus. The correlation between concentration and mean response was significantly correlated for most individual preparations (0.67 to 0.97, mean of r = 0.87 ± 0.08, p < 0.05 in 18/32 preparation-odor pairings). Therefore, excitation scales with changes in odor concentration in MTC glomeruli.

### Different categories of MTC outputs are present when measured with different sensors

We tested whether the four general concentration-response categories were present in MTC glomeruli measured using two different genetically encoded calcium indicators, jRCaMP1b and jGCaMP8m (**Supplementary Figure A2**). Each response type was consistently present in all four preparations sampled across a large concentration range in 4 different odors (**Supplementary Figure A3A-B,** 2 preparations tested for each sensor). These results suggest that the presence of diverse concentration-response relationships are unlikely to be the consequence of sensor properties specific to GCaMP6f.

### Response categories are glomerulus and odor-specific

Our model indicates that different linear and non-linear MTC concentration-response relationships are due to the specific combination of the Hill coefficient and half-activation value of the corresponding ORNs (**Figure 1E**). One prediction of this is that the response category assigned to each glomerulus should be odor-specific. We tested this in a subset of preparations in which responses were measured to at least 2 different odors (N = 479 glomerulus-odor pairings in 15 preparation-odor pairings in 6 different mouse preparations; between 2-4 odors were tested per preparation). Activity maps (ΔF/F) illustrate that response patterns were concentration-dependent and odor-specific (**Figure 4A**). The different response categories of each glomerulus were identified using the previously described criteria and were mapped using different colors. For each preparation, the same glomerulus could exhibit the same kind of response, or entirely different categorical responses (**Figure 4B**). The effect of odor changes from each category to another is illustrated as a 2-dimensional histogram and a network graph (**Figure 4C-D**). Changing the odor resulted in the same response category in 38.8% of glomeruli, with the most common occurrence being NR glomeruli remaining as NR (15.6%) (**Figure 4C-D**, NR vs NR). The remainder of the glomeruli transitioned categories, with the most common outcome being that NR transitioned to I (12.5%) (**Figure 4C-D**, NR vs I). Therefore, the proportions of MTC categories observed across the population are a function of odor-driven variability.

**Figure 4:**
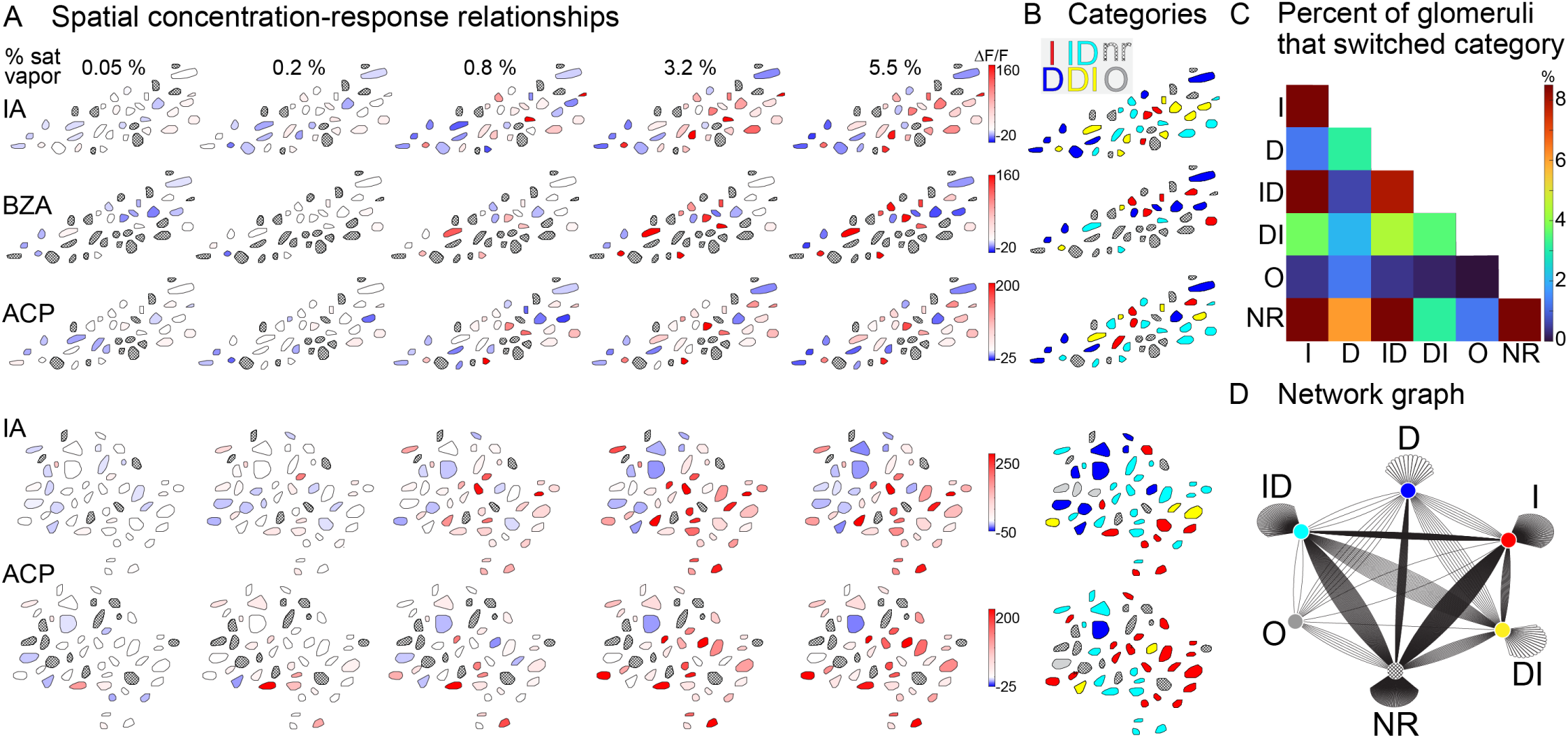
(**A**) Maps of glomerular odor responses (ΔF/F) in different preparations in response to different odors. The color scale is fixed across all concentrations for each preparation-odor pairing. Glomeruli that do not respond to any concentration are indicated with a hatched overlay. (**B**) Maps of glomerular response categories: I, Increasing, D, Decreasing; DI, Decreasing then increasing; ID, Increasing then decreasing, nr, Non-responsive; O, Other type not clearly associated with any category. IA, isoamyl acetate; BZA, benzaldehyde; ACP, acetophenone; MV, methyl valerate. (**C**) A histogram and network graph illustrate how changing the odor changed the proportions of each response type. (**D**) Network graph illustrating transitions between categories as a function of odor change.

### Excitation scales with suppression in MTC glomeruli

Another prediction from our model is that higher levels of excitation should evoke increasing levels of inhibition (**Figure 1B-C**, *dotted line*). We tested this prediction by comparing the mean of all excited glomeruli within a field of view against the mean of all D glomeruli within a field of view. The response to all concentrations was included for each glomerulus if it responded at any concentration. In four exemplar preparation-odor pairings, there was a strong and sometimes statistically significant relationship between normalized excitation and suppression (**Figure 5A-C**, the number of excited and suppressed glomeruli in each preparation-odor pairing is indicated in panel **B**).

**Figure 5:**
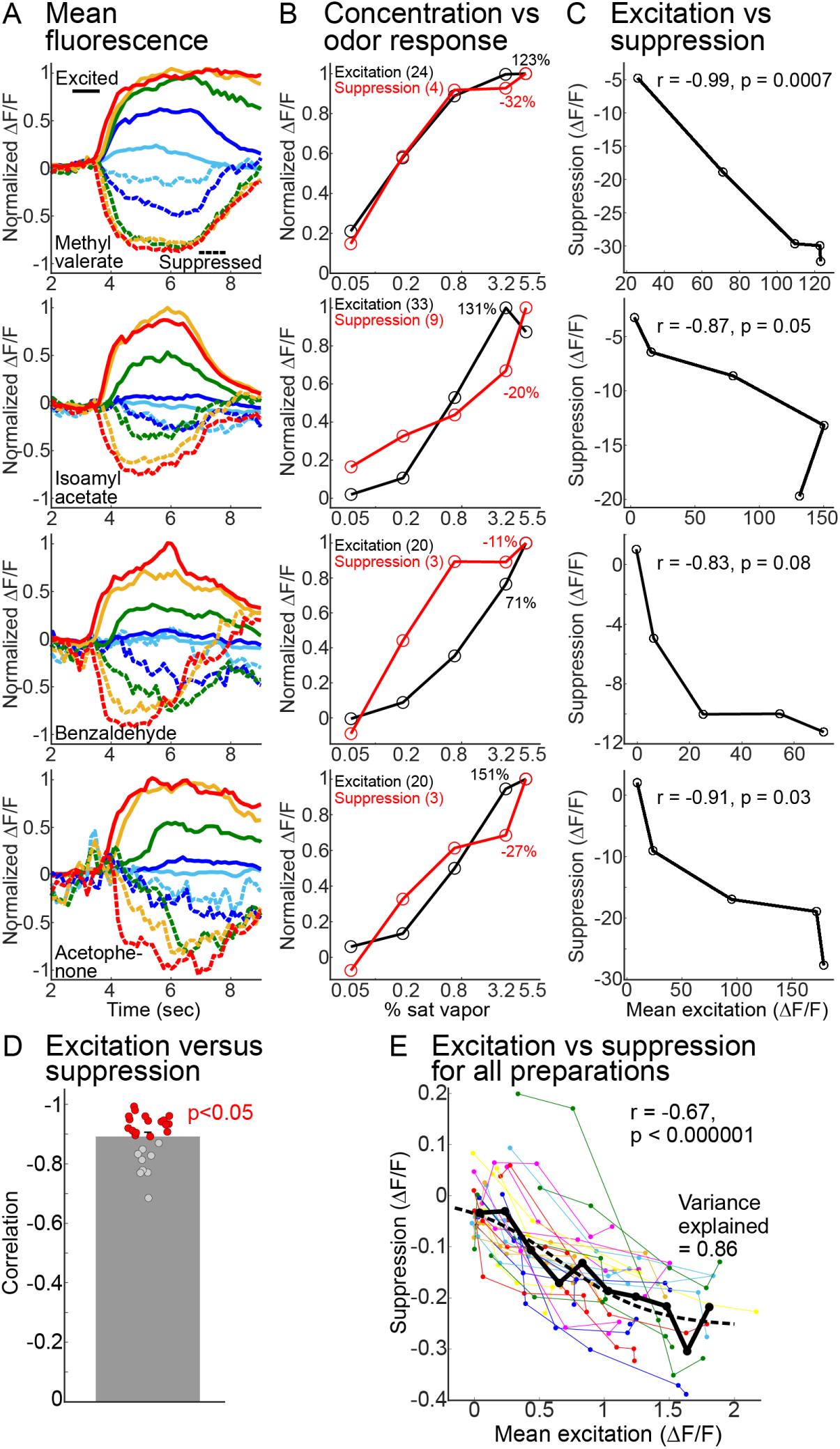
(**A**-**C**) Mean fluorescence time course of excited (solid traces) and suppressed (dashed traces) glomeruli (**A**), concentration versus normalized ΔF/F (**B**), and excitation versus suppression (**C**) for four preparations. The number of glomeruli averaged together and the maximum ΔF/F values are indicated in subpanel **B**. (**D**) Correlation between excitation and suppression for each preparation. (**E**) Mean excitation vs suppression for all preparations. Each individual point is the average of all the excited and suppressed glomeruli within a field of view for one concentration-odor pairing. Measurements from the same preparation-odor pairing are connected with a line. The individual points are binned (solid black line and black circles) and fit to a sigmoid (dashed line).

The relationship between excitation and suppression was significantly correlated in 18 out of the 27 preparation-odor pairings included in the analysis (**Figure 5D**, mean r = 0.89 ± 0.01, statistically significant preparations are indicated in red), and when combining all preparation-odor pairings together (**Figure 5E**, r = -0.67, p < 0.001). Additionally, the individual points from all preparations were binned together, the values of which were fit to a sigmoid (**Figure 5E**, thick black line and circles and the dashed line; different fitting parameters yielded similar fits). Although there was variance in the mean values, there was a clear sigmoidal decline which accounted for a substantial amount of the variance (**Figure 5E**). Therefore, suppression scales with increasing levels of excitation in subsets of MTC glomeruli.

### MTC glomeruli ID concentration-response relationships are heterogeneous and have more sensitive odor responses

Our model predicts that ID responses reflect input from higher affinity ORNs due to receptor saturation, and that differences in the balance of excitation and inhibition should yield differences in the degree of non-monotonicity (**Figure 1D-F**). We quantified the heterogeneity of ID responses using a monotonicity index (MI) which assigns monotonic glomeruli (i.e., I) a value of 0, while non-monotonic glomeruli (e.g., ID) are assigned increasingly negative values depending on the magnitude of the decrease(Escabí *et al*., 2007; Higgins *et al*., 2008).

Concentration-response relationships from simultaneously imaged glomeruli in two different preparations illustrate different I and ID responses (**Figure 6A-B**). We quantified the MI for all responsive glomerulus-odor pairings except for exclusively suppressed glomeruli (D) because they are assigned a strongly negative MI value (**Figure 6C**). Glomeruli binned into different MI ranges illustrate the heterogeneity of ID responses across the MTC population (**Figure 6C**). Within the population of non-suppressed MTC glomeruli, 53% exhibited some degree of non-monotonicity (**Figure 6D**).

**Figure 6:**
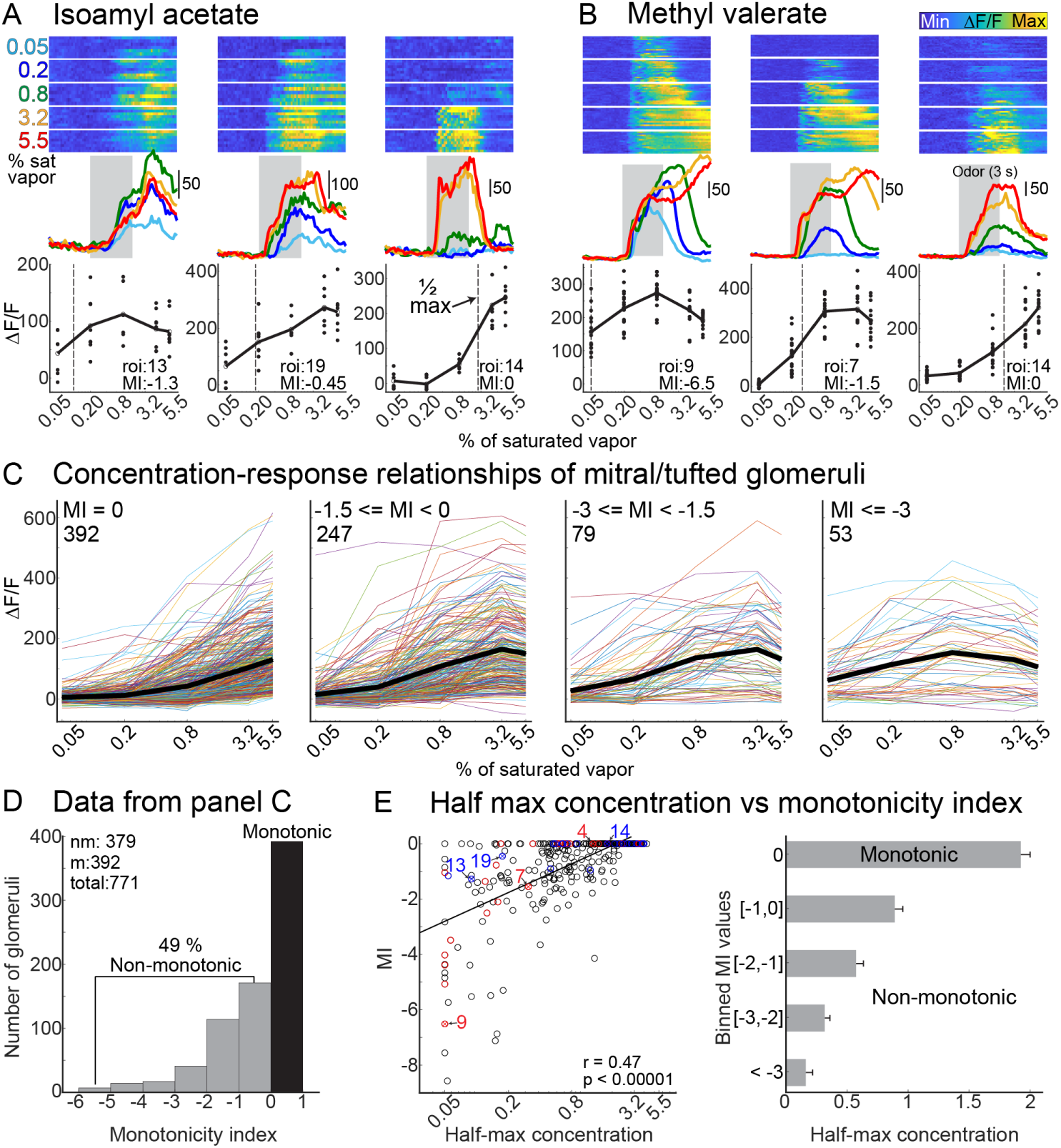
(**A**-**B**) Single trials (top row), mean fluorescence time course (middle) and mean concentration-response relationships (bottom) for three MTC glomeruli in two preparation-odor pairings. The vertical lines in the bottom subpanel of **A**-**B** indicate the half-maximum concentration. (**C**) Concentration response relationships for all responsive MTC glomeruli from all preparations binned by their MI, with the mean indicated with a black line. The number of glomeruli in each bin is in the top left of each panel. (**D**) Histogram of MI values from panel **C**. (**E**) The relationship between half-maximum concentration and MI. The colored circles in the left subpanel indicate measurements from the preparations in panels **A**-**B**.

The MI incorporates response variability at each concentration to assign a value and therefore provides an unbiased approach to defining the two categories (see Eq. 6). Additionally, the MI value was significantly correlated with the linear slope of the concentration-response relationship (r = 0.6, p < 0.0001). Therefore, our metrics accurately capture the differences between monotonic and non-monotonic glomeruli.

We quantified the sensitivity of each MTC glomerulus with piecewise linear interpolation of each concentration-response relationship which was used to quantify the concentration evoking the half maximum response (**Figure 6A-B**, dashed vertical lines). In this analysis, glomeruli in which the response to the lowest concentration was greater than the half-maximum were assigned a value of 0.05% (e.g., **Figure 6B**, e.g., roi 9). ID glomeruli with more negative MI values tended to have lower half-maximum values in our exemplar preparations (**Figure 6A-B**, vertical lines), and across the population (**Figure 6E**, left panel, 324 glomerulus-odor pairings across 17 preparation-odor pairings). This relationship was significantly correlated (r = 0.47, p < 0.0001), and the half-maximum value of I glomeruli was significantly higher than ID glomeruli (**Figure 6E**, right panel, *p <= 0.00046 for all comparisons with MI = 0*). Thus, MTC glomeruli with ID responses tend to reach their half maximum response at lower concentrations than those exhibiting monotonically increasing relationships with odor concentration.

### Imaging the olfactory bulb input-output transformation using dual-color 2-photon imaging

The result that MTC glomeruli with smaller half-maximum values tend to be more non-monotonic is consistent with the model predictions (**Figure 6E**). However, a stronger and more direct test would involve systematically relating odor affinity to glomerular response profiles by functionally imaging ORN and MTC glomeruli. Therefore, we generated a transgenic mouse in which GCaMP6s was expressed in the ORNs, and jRCaMP1b was selectively expressed in MTCs and their apical dendrites innervating the glomerular layer (**Figure 7A-B**). *In vivo* experiments were performed in awake mice in which the dorsal OB was selectively illuminated with 920 nm to measure GCaMP fluorescence, and 1100 nm to measure jRCaMP1b fluorescence (**Figure 7C**, *top row illustrates the mean fluorescence from two different preparations*). In an abundance of caution, we performed all measurements of ORNs and MTCs in separate imaging trials that were illuminated with 920 nm or 1100 nm, respectively, due to the relatively long emission tail present for GCaMP (Akerboom et al., 2012). Imaging trials carried out at 920 nm and 1100 nm are hereafter referred to as ORN and MTC measurements, respectively. Odor responses were measured in 4 different mouse preparations to methyl valerate, isoamyl acetate, acetophenone and methyl salicylate delivered at 6 liquid dilutions ranging from 0.002% to 10% and air (a total of 8 preparation-odor pairings). A frame subtraction analysis revealed that odors evoked changes in glomerular sized regions of interest in both the ORN and MTC measurements (**Figure 7C**, *bottom row*).

**Figure 7:**
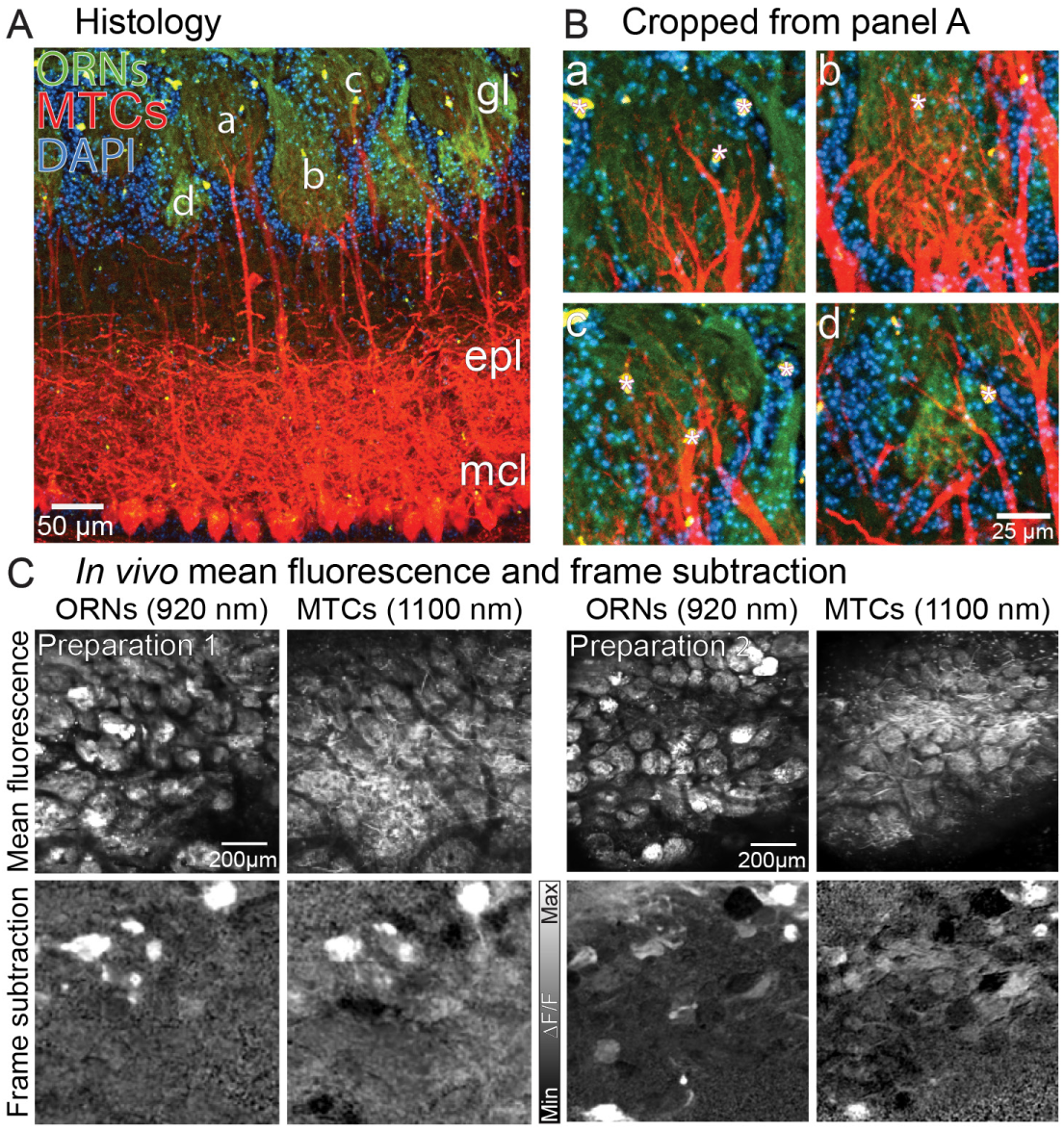
(**A**-**B**) Histology from a transgenic mouse expressing GCaMP6s in the ORNs, and jRCaMP1b in the MTCs. The bright yellow spots indicated with asterisks are lipofuscin autofluorescence. (**C**) Mean fluorescence and frame subtraction odor responses from the ORNs (imaged using 920 nm excitation) and MTCs (imaged using 1100 nm excitation) in 2 mouse preparations.

### Non-monotonic MTC glomerular outputs originate from higher affinity ORN inputs

The ORN and MTC odor response time course and corresponding concentration-response relationships are illustrated for 3 glomeruli from 2 different preparation-odor pairings (**Figure 8A-B**). The most sensitive ORN glomeruli often had correspondingly non-monotonic MTC outputs with small and sometimes negative slopes (**Figure 8A-B**, left most panels). We quantified the affinity of each ORN glomerulus by measuring the half-maximum value of the concentration-response relationship (e.g., **Figure 8A**, *vertical lines in the top row*), and the steepness of each MTC glomerulus by measuring the linear slope of the concentration-response relationship. The relationship between ORN affinity and MTC slope is illustrated for the two preparations in panels **A**-**B** in **Figure 8C** and was significantly correlated when combining all tested glomeruli from 8 preparation-odor pairings (**Figure 8D**, N = 159 glomeruli, glomeruli from different preparation-odor pairings are indicated by color). Finally, this relationship was significantly correlated in 7/8 preparation-odor pairings in the data set (**Figure 8E**).

**Figure 8:**
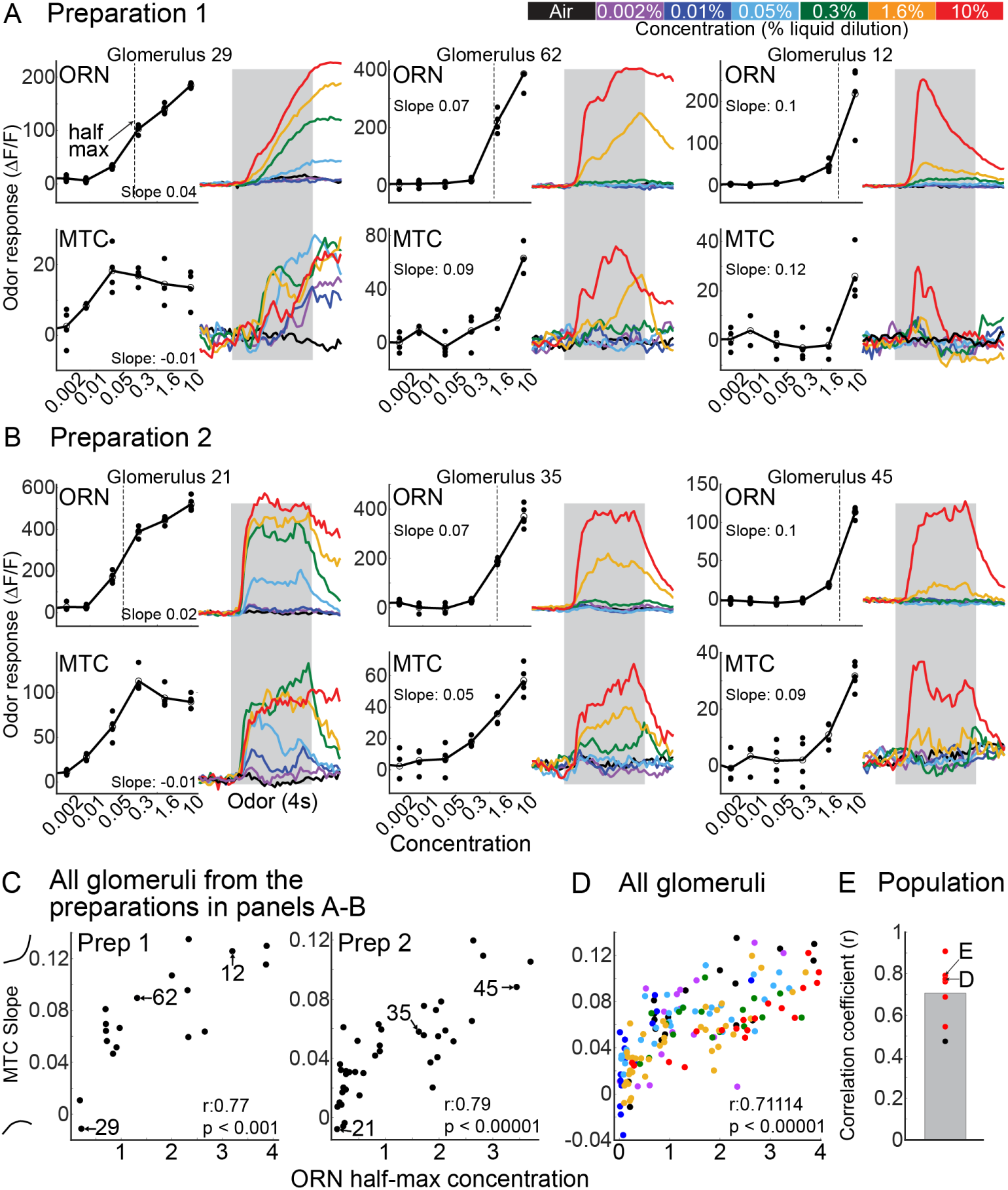
(**A**-**B**) Input-output measurements from three glomeruli from two different mouse preparations. The glomeruli are sorted from lowest (most sensitive) to highest (least sensitive) half-maximum concentration value. (**C**-**D**) The correlation between ORN half-maximum and the slope of the MTC concentration-response relationship for glomeruli in the preparations in panels **A**-**B** (**C**), and for all glomeruli in 8 preparation-odor pairings (**D**). (**E**) Correlation coefficient of each preparation for all 8 preparations.

### Decreasing MTC glomeruli originate from lower sensitivity ORN inputs

Another prediction is that D MTC glomeruli are the consequence of input from ORNs with low sensitivity to an odor ORNs with more sigmoidal concentration-response relationships, in which the rising levels of lateral inhibition exceed the feed-forward ORN excitation. We tested this prediction by measuring the input-output relationship of MTC glomeruli that were clearly suppressed by the odor at the highest concentration (e.g., **Figure 9A**, *output*). Our data set included 64 MTC glomeruli from 5 preparation-odor pairings (3 different animals) that were suppressed by the odor in response to the highest tested concentration with a mean Z-Score change of 5.9 ± 0.4 from baseline (**Figure 9A**, output). Of the MTC glomeruli with suppressed odor responses, 39/64 had an ORN odor responsive to at least one concentration with a minimum change of 5 standard deviations above baseline (e.g., **Figure 9A**, *input*). The odor-responsive inputs to the D MTC glomeruli had a mean half-maximum concentration of 3.4 ± 0.2, which was significantly larger than the half-maximum concentration of all other glomeruli when collapsing across all 5 preparation-odor pairings, or when comparing each preparation-odor pairing individually (**Figure 9B**). Interestingly, the remaining 25/64 D MTC glomeruli originated from ORN inputs that were non-responsive to the same odor (**Figure 9C**, *compare response of MTC and ORN to isoamyl acetate*). In the subset of the non-responsive inputs that were tested to multiple odors, most could be activated by another odor (10/13) (**Figure 9C**, *compare response of ORN to isoamyl acetate and acetophenone or methyl valerate*). This indicates the lack of an ORN response to isoamyl acetate was not due to poor labeling or other methodological explanations. Together these results are consistent with the idea that D glomeruli of MTC are primarily innervated by lower affinity ORN glomeruli, which is consistent with the model predictions.

**Figure 9:**
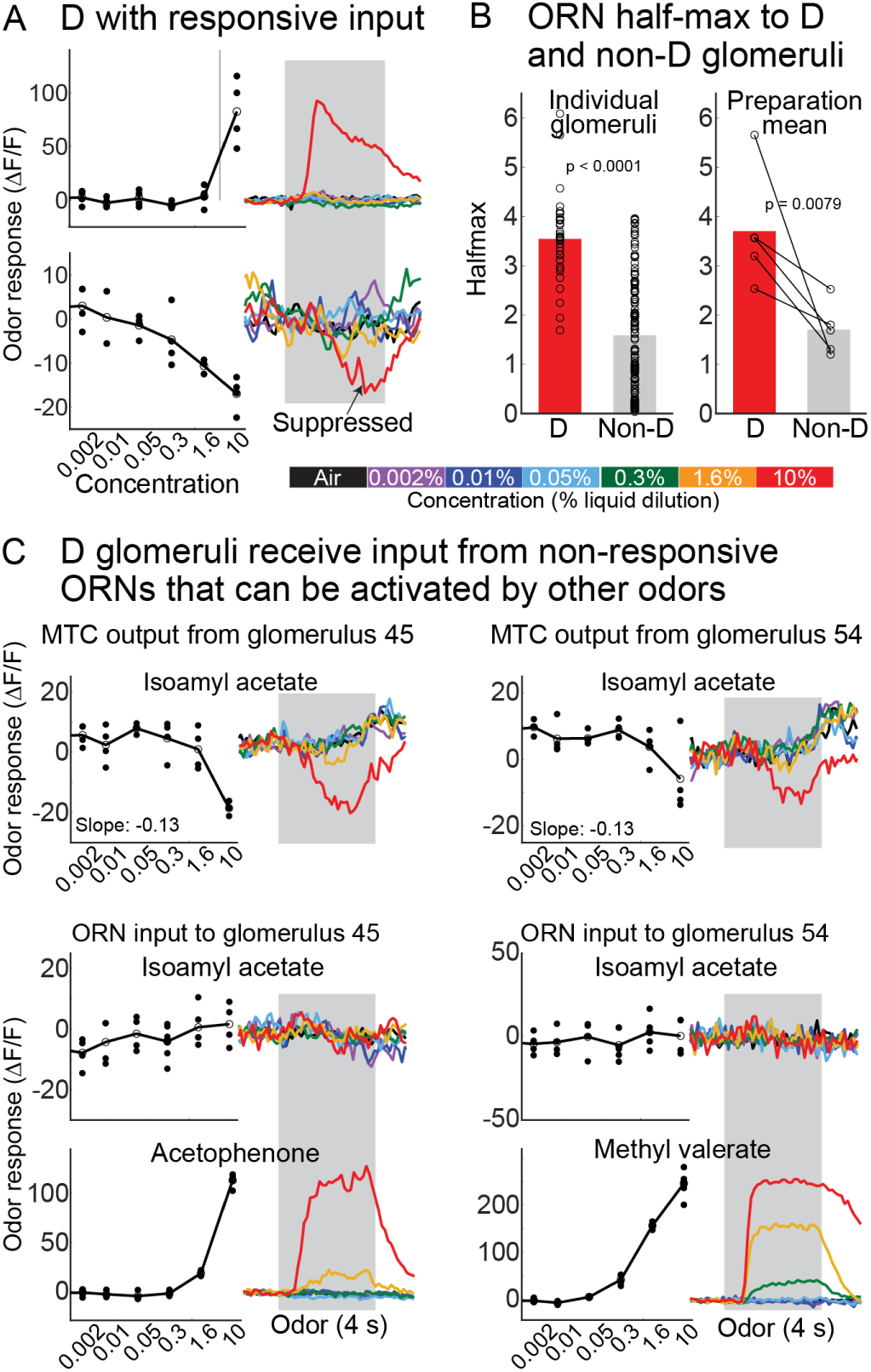
(**A**) An MTC glomerulus with a D concentration-response relationship that receives input from ORN with a high half-maximum concentration. (**B**) Mean half-maximum value of all individual ORN glomeruli with D responses, and for all non-D glomeruli in the same preparation-odor pairings (left) and the mean for each preparation (right). (**C**) T wo MTC glomeruli with D responses that received ORN input that was not responsive to the same odor (isoamyl acetate), but could be activated by other odors (bottom, acetophenone and methyl valerate).

## Discussion

We generated a model of the OB input-output transformation that yielded predictions about how the MTC output of individual glomeruli is shaped by intraglomerular and interglomerular mechanisms. Different ORNs were transformed into different output response types that were determined by the Hill coefficient and half-activation value. Notably, our model predicted the presence of ID responses, which depended on lateral inhibition, were heterogeneous, and originated from ORN input with higher sensitivity to the odor (**Figure 1**). *In vivo* measurements from MTC glomeruli revealed that higher concentrations activated increasing numbers of glomeruli, which exhibited a range of different concentration-response categories that were generally consistent with the model predictions, and which were present when measurements were performed using 3 different optical sensors (**Figure 3** and **Supplementary Figure A3**). Different odors evoked different response categories in 60% of glomerulus-odor pairings, suggesting that the concentration-response function of each MTC glomerulus is driven by network activity (**Figure 4**). Higher concentrations increased the mean level of excitation, which was significantly correlated with the magnitude of suppression present in other glomeruli in the same field of view (**Figure 5**). The model predictions suggest that ID responses are common, heterogeneous, and the consequence of rising levels of lateral inhibition that will begin to suppress sensitive and saturating ORN inputs. We performed dual-color 2-photon Ca^2+^ imaging from ORNs and MTCs in the same glomeruli to confirm this prediction (**Figure 7-8**). Finally, we observed D responses in which glomeruli were exclusively innervated by lower sensitivity or non-responsive ORN inputs, a result that is consistent with rising levels of lateral inhibition (**Figure 9**). The modeling and experimental results indicate that MTC glomerular output is a complex mix of the overall balance of excitation and inhibition innervating that glomerulus.

### Rationale for our modeling choices

A fundamental assumption of the model is that the response function of periglomerular cells (PG) that provide local inhibition is left-shifted relative to that of mitral cells and has a lower saturating level (**Figure 1B**). This provides local half-hat coding, so that weaker input signals are suppressed while stronger ones get through. This is a key element of the model for olfactory coding hypothesized by Cleland (Cleland & Sethupathy, 2006; Cleland, 2010). Although this model has some experimental support, it would be strengthened by future experiments that directly compare the concentration sensitivity of local interneurons with mitral/tufted cells innervating the same glomerulus (e.g., as done for ORNs and MTCs in **Figures 7-9**) (Pinching & Powell, 1971; White, 1972; Gire & Schoppa, 2009; Shao *et al*., 2009; Economo *et al*., 2016).

Another fundamental assumption is that there is lateral inhibition that increases with the odor concentration (e.g., the dashed curve in **Figure 1C**). Without this lateral inhibition the model produces only two response types: I and DI (**Supplemental Figure A1B**). If the half-hat assumption is incorrect, and local interneurons are activated at higher odor concentrations than the mitral cells, then ID mitral cell responses could be produced with local inhibition alone, but in that case DI responses would not be produced through local inhibition. The most parsimonious hypothesis is then that the PG activation function is left-shifted, and primarily responsible for the suppression in DI responses, while lateral inhibition is responsible for the suppression at higher odor concentrations and thereby yielding ID responses.

Our model implements interglomerular lateral inhibition by means of divisive normalization, a well-established mechanism for implementing gain control and contrast enhancement in sensory systems (Carandini & Heeger, 2011). The most widely recognized example is in the visual system, as described by Heeger (1992), where neural responses are modeled as the ratio of input to the summed activity of a pool of neurons (plus a constant). This principle underlies our formulation of normalized ORN activity in Equation (2). Moreover, divisive mechanisms have been observed or proposed in multiple sensory modalities, including olfactory, auditory, and somatosensory systems (Olsen *et al*., 2010; Carandini & Heeger, 2011; Rabinowitz *et al*., 2011). In the olfactory system, interglomerular lateral inhibition is thought to implement a normalization mechanism that can make the glomerular activation pattern more stable across different concentrations (Cleland & Sethupathy, 2006; Cleland *et al*., 2007; Cleland, 2010; Cleland *et al*., 2011). Support for the existence of this mechanism has been shown anatomically and functionally as well (Aungst *et al*., 2003; Olsen & Wilson, 2008; Cleland, 2010; Olsen *et al*., 2010; Zhu *et al*., 2013; Banerjee *et al*., 2015; Storace & Cohen, 2017; Shen *et al*., 2025a). The extent and structure of interglomerular inhibitory coupling is unclear (Banerjee *et al*., 2015; Economo *et al*., 2016; Zavitz *et al*., 2020). We assumed that the lateral inhibition of the input to a glomerulus was from a randomly-selected subpopulation of the glomeruli. The effect of increasing the size of the normalizing subpopulation was to sharpen the boundaries between the different response types (**Figure 1E, F**). It is quite possible that lateral inhibition is more selective than what we assume, for example with preference given to nearby glomeruli. We do not believe that adding specificity would alter the model predictions.

Although we have schematized it as such, the specifics of our model do not require that lateral inhibition function presynaptically at the level of ORNs. The general effect of lateral inhibition in our model only needs to be presynaptic to the mitral cells to function in a similar way. Indeed, recent studies have shown that lateral inhibition can occur presynaptically to mitral cells through an indirect external tufted cell pathway (Banerjee *et al*., 2015). Therefore, it is possible that the “interglomerular” inhibition in our model could reflect a combination of lateral presynaptic inhibition, lateral inhibition occurring at other levels of the bulb including through multi-synaptic pathways presynaptic to the MTCs, through granule cell mediated lateral inhibition, and even feedback from other brain areas (Vucinic *et al*., 2006; Petzold *et al*., 2009; Rothermel *et al*., 2014; Rothermel & Wachowiak, 2014; Banerjee *et al*., 2015; Otazu *et al*., 2015; Kapoor *et al*., 2016; Zak *et al*., 2024).

### Comparison with previous studies

Individual mitral and tufted cells have been reported to exhibit a range of both monotonic and non-monotonic concentration-response relationships (Meredith, 1986; Stopfer *et al*., 2003; Niessing & Friedrich, 2010; Igarashi *et al*., 2012; Kikuta *et al*., 2013; Chae *et al*., 2022; Shen *et al*., 2025b). We propose that differences across studies reporting primarily monotonic versus both monotonic and non-monotonic concentration-response relationships can be explained by our observation that each glomerulus exhibits an odor-specific concentration-response relationship (**Figure 4**) (Kikuta *et al*., 2013; Chae *et al*., 2022). Although mitral cells on average exhibit more non-monotonic relationships than tufted cells, additional studies comparing the activity of individual mitral versus tufted cells innervating the same glomeruli are needed to further understand the relationship of both projection neuron types (Chae *et al*., 2022).

The presence of both linear and non-linear response types is in contrast with two prior studies reporting that MTC glomeruli exhibit exclusively monotonic concentration-response relationships (Fletcher *et al*., 2009; Storace & Cohen, 2017). Because monotonic and non-monotonic response types were often present in the same imaging field of view, the non-monotonic responses cannot be explained as technical limitations associated with our olfactometer, variations in the animal’s respiration, or odor chemistry. In addition, as the model shows, non-monotonic glomerular responses would be expected given what is known about local and interglomerular inhibition (**Figure 1**) (Cleland *et al*., 2011). We propose that these differences primarily reflect the use of epifluorescence versus 2-photon microscopy, which rejects significantly more out-of-focus fluorescence. Epifluorescence measurements from a region of interest will likely reflect a complicated average of the neighboring pixels, which **Figure 4** illustrates can include a combination of I, D, DI and ID glomeruli (Orbach & Cohen, 1983). This is also supported by the observation that MTC glomeruli exhibit less steep concentration-response relationships than their corresponding olfactory receptor neuron input when using epifluorescence imaging (Storace & Cohen, 2017; Storace *et al*., 2019; Leong & Storace, 2024).

The heterogeneity of non-monotonic response types is also different from that described in recent experimental and modeling studies, in which a smaller proportion of measurements exhibited non-monotonic responses (28.7% versus 49%) (Economo *et al*., 2016; Zavitz *et al*., 2020). However, the experimental data in those studies were collected over a much smaller concentration range in anesthetized animals, which may have reduced the ability to detect changes that would occur over a larger range. While the study of Zavitz et al., 2020 captured the monotonic responses reported in (Economo *et al*., 2016), they did not account for the non-monotonic responses that were so frequent in our data.

Current models of how inhibitory circuits are shaped by excitation propose that inhibition either increases broadly across the glomerular population in a manner that scales with the magnitude of excitatory input, or increases selectively (Cleland *et al*., 2007; Banerjee *et al*., 2015; Economo *et al*., 2016; Zavitz *et al*., 2020). ID responses have been rarely described at the level of the input to the olfactory bulb (Wachowiak & Cohen, 2001; Bozza *et al*., 2004; Carey *et al*., 2009; Lecoq *et al*., 2009; Storace & Cohen, 2017; Inagaki *et al*., 2020; Zak *et al*., 2020; Martelli & Storace, 2021; Platisa *et al*., 2022). Our theoretical and experimental results suggest that ID glomeruli are likely to reflect processing that occurs within the OB as higher sensitivity ORN input saturates and is weakened due to rising lateral inhibition (**Figures 1**, **6** and **8**). The result that higher levels of suppression are significantly correlated with increasing excitation is consistent with studies indicating that the responses of interneurons that mediate lateral connectivity exhibit activity that scales with increasing odor concentrations (Zhu *et al*., 2013; Banerjee *et al*., 2015; Storace *et al*., 2019).

However, another study reported that the amount of excitation in MTC glomeruli was only weakly predictive of the magnitude of suppression (Economo *et al*., 2016). We propose that a few methodological differences can explain the different conclusions. Our study sampled each glomerular field of view across a large range of odor concentrations, which increased the likelihood that we would observe glomeruli exhibiting this relationship. Additionally, glomeruli that were suppressed at the highest concentration were selected for analysis and their responses to each concentration were analyzed even if they were non-responsive at lower concentrations. This facilitated the identification of glomeruli with graded transitions from non-responsive to suppressed. Finally, we excluded glomeruli that we categorized as DI, which typically responded with suppression to lower concentrations, and transitioned to excitation at higher concentrations.

Our finding that more than half of the MTC glomeruli exhibit monotonic concentration-response relationships suggests it is ubiquitous. We also found that most MTC glomeruli exhibiting monotonic concentration-response relationships were well fit by the Hill equation and have Hill coefficients consistent with previous ORN measurements (Wachowiak & Cohen, 2001; Storace & Cohen, 2017; Zak *et al*., 2020; Platisa *et al*., 2022). This does not mean that these MTC glomeruli are not affected by inhibitory input. In fact, in the model all glomeruli are subject to both local and lateral inhibition, but in many cases the concentration-response function is monotonically increasing (**Figure 1D**, I). Future studies with a more comprehensive analysis of the olfactory bulb input-output transformation alters concentration-response relationships are required to resolve the question of how each individual glomerulus is distinctly impacted by OB processing.

ORNs can undergo rapid adaptation in response to prolonged or high frequency sampling of an odor stimulus (Duchamp-Viret *et al*., 2000; Verhagen *et al*., 2007; Lecoq *et al*., 2009). It remains unclear how this kind of adaptation influences the glomerular output response, and whether it plays a role in the different response types we describe. In our experiments in which we measured both the ORNs and MTCs, we did not observe rapid adaptation in the ORN input to MTC glomeruli with ID response types (**Figure 8A**, glomerulus 29; **Figure 8B**, glomerulus 21). Additionally, the result that the example ID MTC glomeruli in **Figure 3A**, **Figure 3B**, **Figure 6A** and **Figure 6B** ID glomerulus remained elevated throughout the odor duration made it unlikely to be the sole mechanism for ID response types. However, because we did not measure the ORN input to all glomeruli in our data set, it is impossible to rule out the possible contribution of ORN adaptation in some non-monotonic responses. A more comprehensive data set including more input-output measurements are needed to understand how ORN adaptation transforms the olfactory bulb input-output transformation.

### Methodological considerations

Because action potentials trigger a rapid influx of calcium into neurons, calcium indicators provide an indirect report of neural activity. Additionally, calcium sensors each have distinct biophysical properties, which include their affinity for calcium, the slope of this binding relationship, and the speed at which they can transform calcium changes into changes in fluorescence. For example, GCaMP6f, jGCaMP8m and jRCaMP1b have calcium affinities of 375 nM, 712 nM, and 108 nM, and Hill coefficients of 2.27, 1.6 and 1.92, respectively (Chen *et al*., 2013; Badura *et al*., 2014; Dana *et al*., 2016; Helassa *et al*., 2016; Zhang *et al*., 2023).

Despite their non-linear Hill coefficients, there is a relatively strong relationship between spiking activity and changes in fluorescence. GCaMP3 and GCaMP5G exhibit continuous increases in ΔF/F to electrical stimulation in cultured neurons and in response to spiking activity in olfactory bulb mitral cells (Tian *et al*., 2009; Akerboom *et al*., 2012; Kato *et al*., 2012; Wachowiak *et al*., 2013), and similar observations have been made for GCaMP6f, jRCaMP1b and jGCaMP8 (Chen *et al*., 2013; Dana *et al*., 2016; Zhang *et al*., 2023). However, the ability to detect neural activity across a large range of spiking is ultimately limited by the dynamic range of the protein sensor itself (Wachowiak *et al*., 2013).. Because mitral/tufted cells have relatively high spontaneous firing rates, the ability of GECIs to report the full range of their spiking activity may be limited (Rinberg *et al*., 2006; Zhang *et al*., 2023).

The observation that ID and DI responses were present in multiple data sets using three different calcium indicators with different biophysical properties suggests that they cannot be easily explained as limitations of the indicator (**Figure 3**, **Figure 8**, **Supplementary Figure A3**). Regardless, future experiments are needed in which input-output measurements are performed using spectrally distinct voltage sensors expressed in the ORNs and MTCs (Storace & Cohen, 2017; Kannan *et al*., 2018; Platisa *et al*., 2022; Leong & Storace, 2024).

Mitral cell action potentials initiated at the soma backpropagate into primary dendrites (Bischofberger & Jonas, 1997; Chen *et al*., 1997; Chen *et al*., 2002; Christie & Westbrook, 2003; Djurisic *et al*., 2004), which evoke corresponding calcium transients (Charpak *et al*., 2001; Debarbieux *et al*., 2003; Wachowiak *et al*., 2013). Action potential driven calcium transients in the apical dendrites are substantially larger than those evoked by pre-synaptic input (Charpak *et al*., 2001; Zhou *et al*., 2006; Kato *et al*., 2012). Therefore, although MTC glomerular signals do not exclusively reflect somatic spiking activity, their measurements are a reasonable way to measure changes across the glomerular input-output transformation.

The Tbx21-cre transgenic strain used in our study drives expression in both mitral and tufted cells (Mitsui *et al*., 2011; Storace & Cohen, 2021), which can have distinct concentration-response relationships and are differently embedded into the glomerular network (Nagayama *et al*., 2004; Fukunaga *et al*., 2012; Igarashi *et al*., 2012; Chae *et al*., 2022; Shen *et al*., 2025b). Therefore, our 2-photon imaging from the apical dendrites of MTCs innervating glomeruli necessarily include signals from both cell types. Understanding the distinct role(s) of mitral versus tufted cells in glomerular signaling require future experiments incorporating more selective targeting to the two cell populations (Rothermel *et al*., 2013; Wachowiak *et al*., 2013; Koldaeva *et al*., 2021).

Here we delivered changes in odor concentration using two different olfactometer systems, one in which pure odor was delivered across a 100-fold range of percent of saturated vapor, and the other in which odors were delivered at a constant 10% of saturated vapor but were diluted in mineral oil across a much larger range. Although DI response types could be detected at our second lowest concentration (0.2 % sat vapor), the non-monotonic component of ID glomeruli, and many detectable D responses typically emerged at or above 3.2 % of saturated vapor (**Figure 6**), which is a concentration range that an animal is unlikely to experience in its natural environment (Wachowiak *et al*., 2025). We propose that the ID responses we describe are more likely to occur in more natural contexts which can contain multiple odor sources that will activate multiple glomeruli and therefore higher levels of lateral inhibition.

### Functional relevance of non-monotonic concentration-response relationships

Non-monotonic intensity relationships (“ID” in our study) have been described in other sensory systems as a mechanism by which the brain can encode absolute or relative intensity differences (Sutter & Loftus, 2003; Wehr & Zador, 2003; Polley *et al*., 2007; Higgins *et al*., 2010). In the auditory cortex, non-monotonic neurons exhibit unique characteristics including being particularly sensitive to behavioral training and exhibiting unique adaptive properties (Polley *et al*., 2004; Polley *et al*., 2006; Watkins & Barbour, 2008). Although our model indicates that these glomeruli are a natural consequence of the interplay between intra- and interglomerular inhibition, it remains to be tested whether these strongly suppressed glomeruli are functionally unique.

One advantage of having four distinct response types to an odor over a range of concentrations is that it provides a broader range of coverage of the MTC state space. As a simple example we consider only three glomeruli, with output MTC_1_, MTC_2_, and MTC_3_. The response amplitude of each to an odor at a given concentration would be a point in the MTC_1_-MTC_2_-MTC_3_ state space, and the response to that odor over a range of concentrations would be a set of points that can be thought of as a directed curve in that space, which we refer to as a trajectory (**Figure 10**). The neural coding of an odor across concentrations in the olfactory bulb would then be a trajectory in the MTC state space. The response to a second odor over a similar range of concentrations would produce a different trajectory. In both cases, the trajectory should be considered a cloud of points that reflect variations in the fluctuations of molecular components present in natural odor stimuli (whether due to differences in chemistry or environmental interactions). An animal would learn to recognize an odor in a concentration invariant manner as the population activation along that trajectory based on experience. To illustrate how different response types impact coverage of the MTC state space we show a collection of trajectories all having an I response to an odor (black curves in **Figure 10**) and an equal number having a mix of monotonic and non-monotonic response types (red curves in **Figure 10**). Each trajectory is based on calcium measurements from 3 glomeruli. When the responses are all the I type they cover only a small region of the state space. In contrast, when non-monotonic responses are included, the coverage is much more extensive. The greater coverage makes it easier to discriminate one odor from another, and even though the dimension of the actual MTC state space is much larger than 3 (equal to the number of different receptor types), it is still advantageous to increase the coverage of that space, as is done by having different response types to odors.

**Figure 10:**
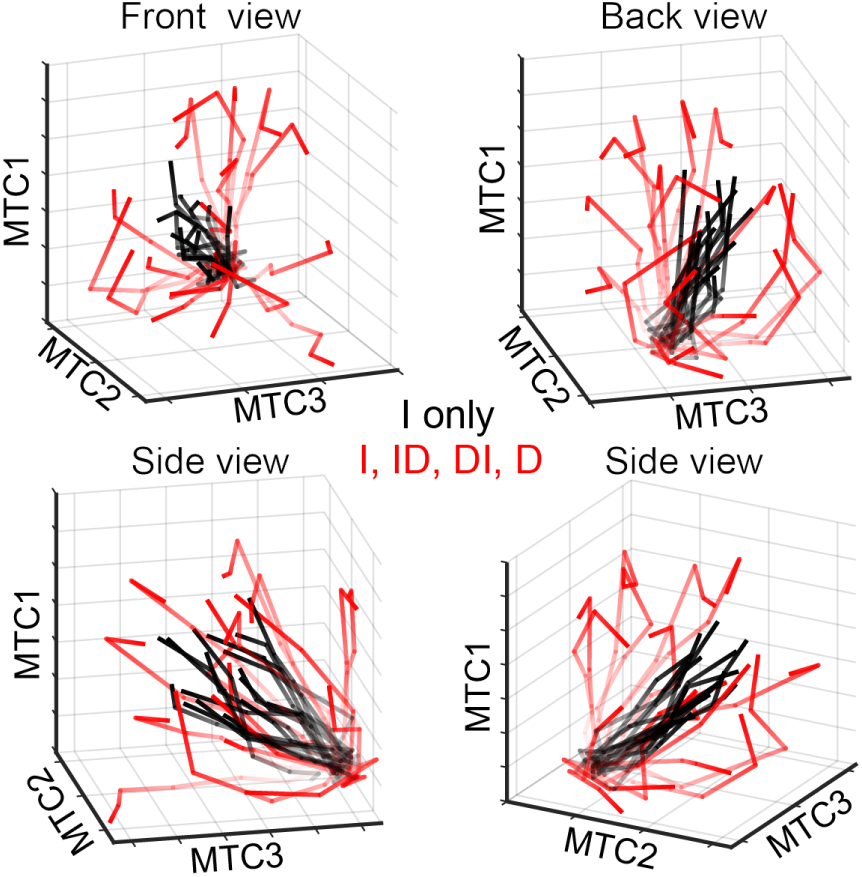
Model of how the different MTC response types influence odor coding. Each line in this model illustrates the response of 3 MTC glomeruli to a single odor across concentration changes. Concentration changes are represented as the changing transparency of each line. MTC glomeruli with only increasing responses tended to cluster together in space (black lines). In contrast, MTC glomeruli with different response categories had response trajectories that diverged from one another (red lines). Each panel illustrates the projections from a different angle.

### Future studies

It is necessary to know the concentration-response relationships of the input to each glomerulus to further understand the transformation occurring within each glomerulus. Knowing the input-output relationship will allow for additional tests of our proposed model, including a more precise test of the relationship between affinity and ID responses (**Figure 1**). Importantly, the model predicted that DI responses could be due to intraglomerular processing, while ID responses required interglomerular processing. Future studies are needed to test this hypothesis by measuring the concentration-response relationships in response to selective manipulation of different cell types or more precise control over the input stimulus (Smear *et al*., 2013; Banerjee *et al*., 2015; Braubach *et al*., 2018; Burton *et al*., 2022).

Here we tested several predictions involving inhibition and the role of the half-activation value (**Figure 1E**, **Figures 4-9**). However, our model indicates that the Hill coefficient also plays an important role in the transformation of each glomerulus. For example, ORN responses with a half-maximum of ∼0.4 value transition from primarily I response types to ID depending on the Hill coefficient. However, accurate Hill fits require measuring the response at the saturation concentration, and our limited data included many ORNs that did not saturate (e.g., **Figure 8A**, glomerulus 62). Future studies involving a more comprehensive input-output comparison can perform a more thorough test of the impact of the interaction between the half-activation and Hill coefficient of ORNs on MTC glomeruli. Importantly, our half-activation values of the least sensitive ORN glomeruli are likely underestimates, which would not alter the conclusions of our study (e.g., **Figure 8A**, glomeruli 62 and 12).

Here we focus on the amplitude of the calcium response rather than the timing, and our mathematical model likewise is independent of response timing. It is likely, however, that timing is also important for the neural coding of odor (Shusterman *et al*., 2011; Smear *et al*., 2011; Chong *et al*., 2019; Ackels *et al*., 2021). Indeed, the “primacy coding” scheme hypothesizes that only those receptors that act earliest are responsible for the neural coding of an odor (Wilson *et al*., 2017). In this model, whether the MTC response is monotonic or non-monotonic is of little relevance since what matters is the timing of the response. This coding strategy greatly limits the potential that is available through the variety of amplitude responses (I, D, DI, ID) produced at different odor concentrations that we and others have found. Both timing and amplitude provide useful information that can be employed in the neural coding of odor over different concentrations. Temporal dynamics of glomerular responses to an odor could be incorporated into the model by using time-dependent differential equations at the level of ORNs and MTCs.

The present study used monomolecular odors, yet natural odors are often mixtures of different components that can interact antagonistically at the ORN receptor level (Inagaki *et al*., 2020; Xu *et al*., 2020; Zak *et al*., 2020). An odor mixture that causes antagonism in a particular OR type will necessarily right shift its ORN binding curve to an odor (i.e., make it less sensitive). At the same time, the mixture will increase the magnitude of lateral inhibition by activating other OR types (**Figure 1C**). One prediction from our model is therefore that antagonized ORNs will result in MTCs outputs that shift towards the D and DI categories (moving downward in **Figure 1E**). Thus, we suggest that antagonism or mixtures need not be thought of as interfering with the signal that an odorant sends to a particular MTC but instead changes the way that the MTC codes for that odor.

The model can also be further developed to consider the impact of different interglomerular inhibitory network structures (Zavitz *et al*., 2020). Spatial effects may be heterogeneous, for example, and there could be spatial or functional clustering, so that the lateral inhibition targeted to a glomerulus could be preferentially from nearby glomeruli, or from glomeruli responsive to very distinct odors.

### Conclusions

Our comparisons of input and output and mathematical modeling provide direct evidence of a functional transformation occurring in the OB. Non-monotonic concentration response relationships in MTCs are common and expected given the ways that local and lateral inhibition can shape MTC activity. This transformation is likely to play a key role in facilitating odor discrimination and the ability for an animal to achieve perceptual concentration invariance.

## Additional Information

### Data availability

All data are available to The Journal of Physiology and interested parties upon reasonable request.

### Author contributions

D.A.S. and R.B. were involved in the conception or design of the work. All authors were involved in the acquisition, analysis or interpretation of data for the work. D.A.S. wrote the first draft of the manuscript, and all authors were involved in revising it critically for important intellectual content. All authors approved the final version of the manuscript, agree to be accountable for all aspects of the work in ensuring that questions related to the accuracy or integrity of any part of the work are appropriate investigated and resolved; and all persons designed as authors qualify for authorship, and all those who quality for authorship are listed.

## Acknowledgements

We dedicate this paper to our mentor Lawrence B. Cohen who always knew what to do. Thanks to Dr. Evan Lloyd and members of the Storace laboratory for thoughtful discussion of the manuscript.

## Appendix

**Supplementary Figure A1:**
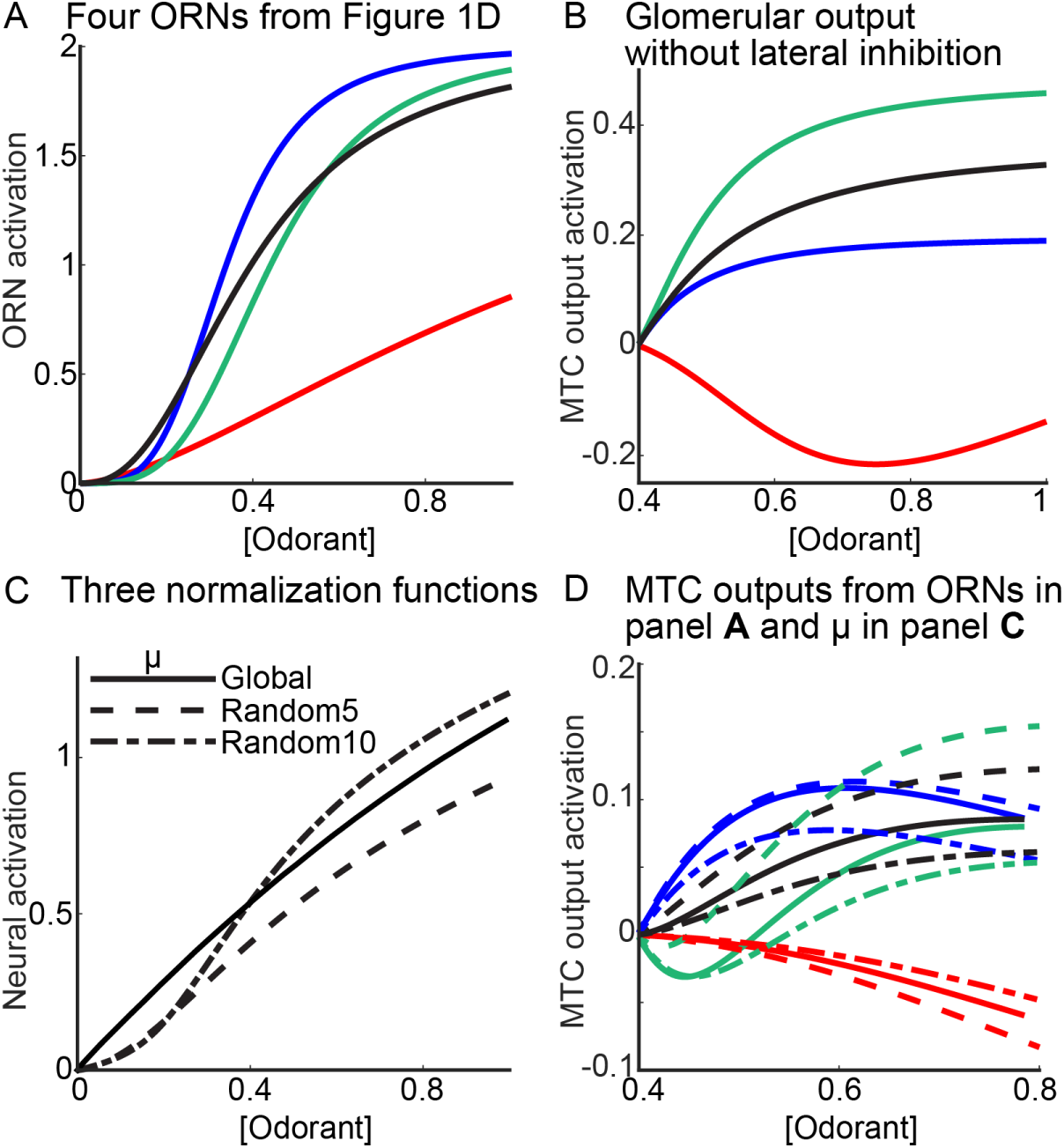
Input and output curves from the mathematical model. (**A**) The 4 exemplar ORN concentration-response functions from Figure 1D. (**B**) The 4 inputs from panel **A** have been processed by the local inhibition circuit (only), resulting in three I and one DI MTC response functions. Importantly, no ID responses were generated in the absence of lateral inhibition. (**C**) Three different ORN functions used to implemental interglomerular processing. The solid “μ_global_” distribution is identical to the black dashed line in Figure 1C. Randomly sampling and then averaging 5 and 10 signals from that figure’s ORN population yielded the “μ_Random5_” and “μ_Random10_” distributions, respectively (*different dashed lines*). (**D**) The 4 inputs from panel **A** processed using local inhibition and lateral inhibition implemented via the different functions in panel **C**. All 4 MTC categories were present irrespective of which population of ORNs were selected for interglomerular processing.

**Supplementary Figure A2:**
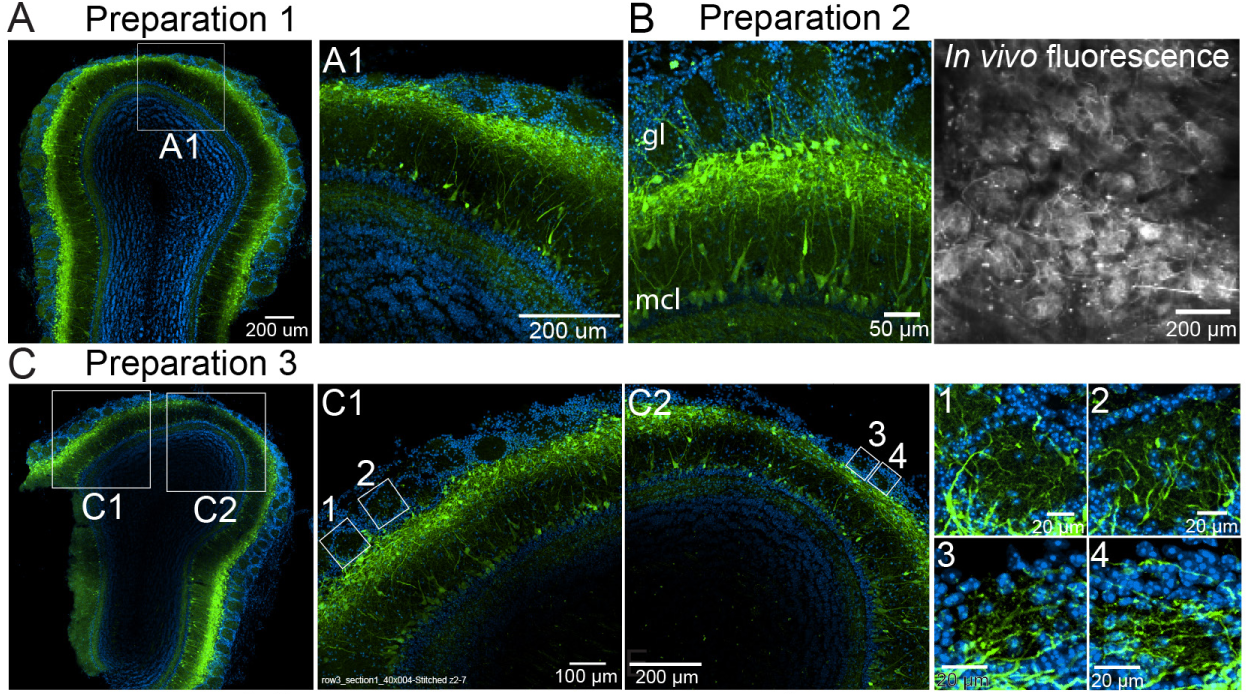
jGCaMP8m expression in the OB in a cross between the Tbx21-Cre and the TIGRE2-jGCaMP8m transgenic lines. (**A-C**) Images from three different mouse preparations. Panel **B** includes *in vivo* 2-photon fluorescence from the glomerular layer from that preparation. Panel **C** includes cropped images from the glomerular layer with an adjusted color lookup table. gl, glomerular layer; mcl, mitral cell layer.

**Supplementary Figure A3:**
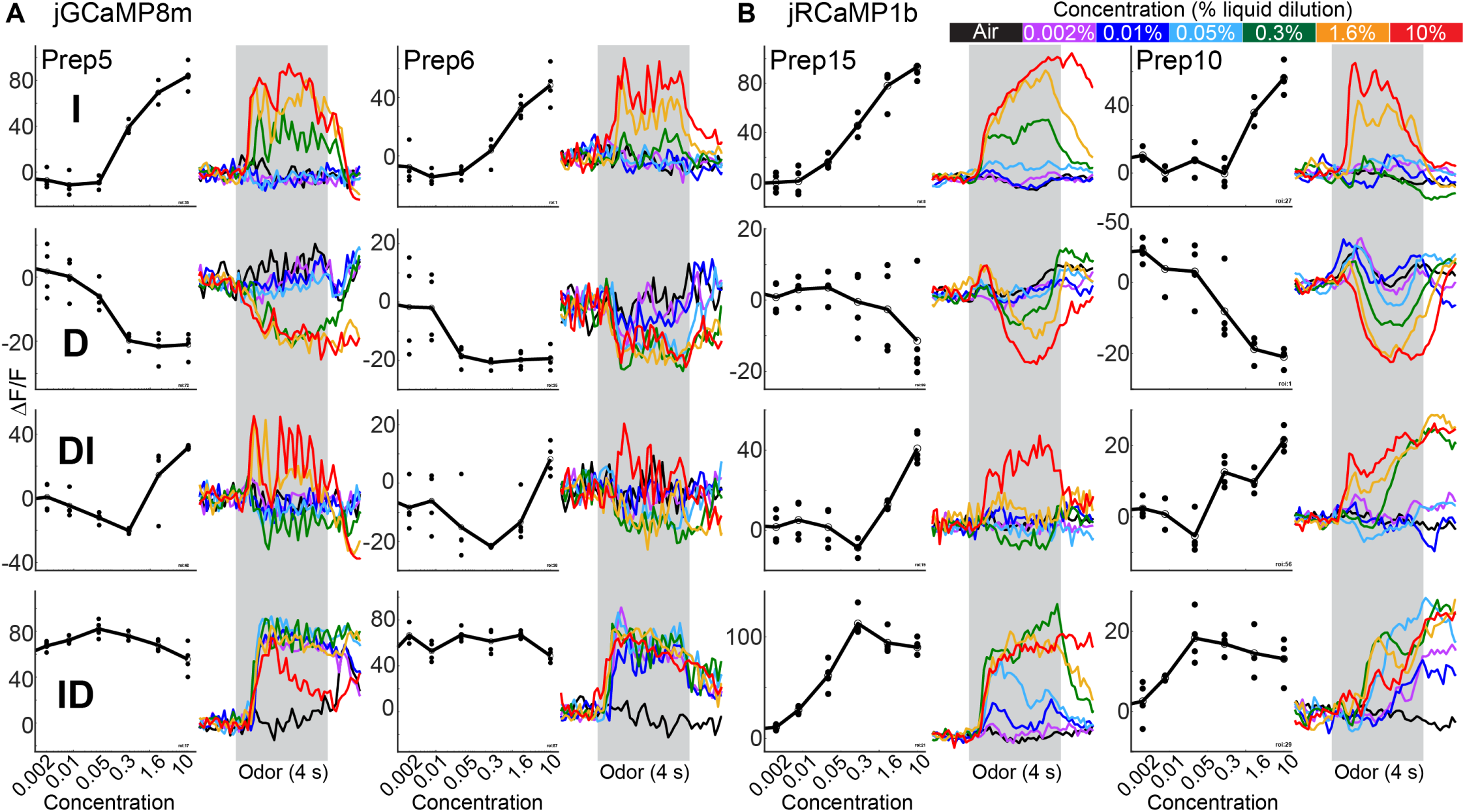
MTC glomerular responses with increasing, decreasing, decreasing-increasing and increasing-decreasing response categories measured with jGCaMP8m (**A**) and jRCaMP1b (**B**).

## Bibliography

Ackels T, Erskine A, Dasgupta D, Marin AC, Warner TPA, Tootoonian S, Fukunaga I, Harris JJ & Schaefer AT. (2021). Fast odour dynamics are encoded in the olfactory system and guide behaviour. Nature 593, 558–563.

Akerboom J, Chen TW, Wardill TJ, Tian L, Marvin JS, Mutlu S, Calderon NC, Esposti F, Borghuis BG, Sun XR, Gordus A, Orger MB, Portugues R, Engert F, Macklin JJ, Filosa A, Aggarwal A, Kerr RA, Takagi R, Kracun S, Shigetomi E, Khakh BS, Baier H, Lagnado L, Wang SS, Bargmann CI, Kimmel BE, Jayaraman V, Svoboda K, Kim DS, Schreiter ER & Looger LL. (2012). Optimization of a GCaMP calcium indicator for neural activity imaging. J Neurosci 32, 13819–13840.

Araneda RC, Kini AD & Firestein S. (2000). The molecular receptive range of an odorant receptor. Nat Neurosci 3, 1248–1255.

Aungst JL, Heyward PM, Puche AC, Karnup SV, Hayar A, Szabo G & Shipley MT. (2003). Centre-surround inhibition among olfactory bulb glomeruli. Nature 426, 623–629.

Badura A, Sun XR, Giovannucci A, Lynch LA & Wang SS. (2014). Fast calcium sensor proteins for monitoring neural activity. Neurophotonics 1, 025008.

Banerjee A, Marbach F, Anselmi F, Koh MS, Davis MB, Garcia da Silva P, Delevich K, Oyibo HK, Gupta P, Li B & Albeanu DF. (2015). An Interglomerular Circuit Gates Glomerular Output and Implements Gain Control in the Mouse Olfactory Bulb. Neuron 87, 193–207.

Bischofberger J & Jonas P. (1997). Action potential propagation into the presynaptic dendrites of rat mitral cells. J Physiol 504 (Pt 2), 359–365.

Bozza T, McGann JP, Mombaerts P & Wachowiak M. (2004). In vivo imaging of neuronal activity by targeted expression of a genetically encoded probe in the mouse. Neuron 42, 9–21.

Braubach O, Tombaz T, Geiller T, Homma R, Bozza T, Cohen LB & Choi Y. (2018). Sparsened neuronal activity in an optogenetically activated olfactory glomerulus. Sci Rep 8, 14955.

Buck L & Axel R. (1991). A novel multigene family may encode odorant receptors: a molecular basis for odor recognition. Cell 65, 175–187.

Burton SD, Brown A, Eiting TP, Youngstrom IA, Rust TC, Schmuker M & Wachowiak M. (2022). Mapping odorant sensitivities reveals a sparse but structured representation of olfactory chemical space by sensory input to the mouse olfactory bulb. Elife 11.

Carandini M & Heeger DJ. (2011). Normalization as a canonical neural computation. Nat Rev Neurosci 13, 51–62.

Carey RM, Verhagen JV, Wesson DW, Pirez N & Wachowiak M. (2009). Temporal structure of receptor neuron input to the olfactory bulb imaged in behaving rats. J Neurophysiol 101, 1073–1088.

Chae H, Banerjee A, Dussauze M & Albeanu DF. (2022). Long-range functional loops in the mouse olfactory system and their roles in computing odor identity. Neuron 110, 3970–3985.e3977.

Charpak S, Mertz J, Beaurepaire E, Moreaux L & Delaney K. (2001). Odor-evoked calcium signals in dendrites of rat mitral cells. Proc Natl Acad Sci U S A 98, 1230–1234.

Chen TW, Wardill TJ, Sun Y, Pulver SR, Renninger SL, Baohan A, Schreiter ER, Kerr RA, Orger MB, Jayaraman V, Looger LL, Svoboda K & Kim DS. (2013). Ultrasensitive fluorescent proteins for imaging neuronal activity. Nature 499, 295–300.

Chen WR, Midtgaard J & Shepherd GM. (1997). Forward and backward propagation of dendritic impulses and their synaptic control in mitral cells. Science 278, 463–467.

Chen WR, Shen GY, Shepherd GM, Hines ML & Midtgaard J. (2002). Multiple modes of action potential initiation and propagation in mitral cell primary dendrite. J Neurophysiol 88, 2755–2764.

Chen Y, Chen X, Baserdem B, Zhan H, Li Y, Davis MB, Kebschull JM, Zador AM, Koulakov AA & Albeanu DF. (2022). High-throughput sequencing of single neuron projections reveals spatial organization in the olfactory cortex. Cell 185, 4117–4134.e4128.

Chong E, Moroni M, Wilson C, Shoham S, Panzeri S & Rinbert D. (2019). Manipulating synthetic optogenetic odors reveals the coding logic of olfactory perception. BioRXiv.

Christie JM & Westbrook GL. (2003). Regulation of backpropagating action potentials in mitral cell lateral dendrites by A-type potassium currents. J Neurophysiol 89, 2466–2472.

Cleland TA. (2010). Early transformations in odor representation. Trends Neurosci 33, 130–139.

Cleland TA, Chen SY, Hozer KW, Ukatu HN, Wong KJ & Zheng F. (2011). Sequential mechanisms underlying concentration invariance in biological olfaction. Front Neuroeng 4, 21.

Cleland TA, Johnson BA, Leon M & Linster C. (2007). Relational representation in the olfactory system. Proc Natl Acad Sci U S A 104, 1953–1958.

Cleland TA & Sethupathy P. (2006). Non-topographical contrast enhancement in the olfactory bulb. BMC Neurosci 7, 7.

Daigle TL, Madisen L, Hage TA, Valley MT, Knoblich U, Larsen RS, Takeno MM, Huang L, Gu H, Larsen R, Mills M, Bosma-Moody A, Siverts LA, Walker M, Graybuck LT, Yao Z, Fong O, Nguyen TN, Garren E, Lenz GH, Chavarha M, Pendergraft J, Harrington J, Hirokawa KE, Harris JA, Nicovich PR, McGraw MJ, Ollerenshaw DR, Smith KA, Baker CA, Ting JT, Sunkin SM, Lecoq J, Lin MZ, Boyden ES, Murphy GJ, da Costa NM, Waters J, Li L, Tasic B & Zeng H. (2018). A Suite of Transgenic Driver and Reporter Mouse Lines with Enhanced Brain-Cell-Type Targeting and Functionality. Cell 174, 465-480 e422.

Dana H, Mohar B, Sun Y, Narayan S, Gordus A, Hasseman JP, Tsegaye G, Holt GT, Hu A, Walpita D, Patel R, Macklin JJ, Bargmann CI, Ahrens MB, Schreiter ER, Jayaraman V, Looger LL, Svoboda K & Kim DS. (2016). Sensitive red protein calcium indicators for imaging neural activity. Elife 5.

Debarbieux F, Audinat E & Charpak S. (2003). Action potential propagation in dendrites of rat mitral cells in vivo. J Neurosci 23, 5553–5560.

Djurisic M, Antic S, Chen WR & Zecevic D. (2004). Voltage imaging from dendrites of mitral cells: EPSP attenuation and spike trigger zones. J Neurosci 24, 6703–6714.

Duchamp-Viret P, Duchamp A & Chaput MA. (2000). Peripheral odor coding in the rat and frog: quality and intensity specification. J Neurosci 20, 2383–2390.

Economo MN, Hansen KR & Wachowiak M. (2016). Control of Mitral/Tufted Cell Output by Selective Inhibition among Olfactory Bulb Glomeruli. Neuron 91, 397–411.

Escabí MA, Higgins NC, Galaburda AM, Rosen GD & Read HL. (2007). Early cortical damage in rat somatosensory cortex alters acoustic feature representation in primary auditory cortex. Neuroscience 150, 970–983.

Fantana AL, Soucy ER & Meister M. (2008). Rat olfactory bulb mitral cells receive sparse glomerular inputs. Neuron 59, 802–814.

Firestein S & Zufall F. (1993). Membrane currents and mechanisms of olfactory transduction. Ciba Found Symp 179, 115–126; discussion 126-130, 147-119.

Fletcher ML, Masurkar AV, Xing J, Imamura F, Xiong W, Nagayama S, Mutoh H, Greer CA, Knopfel T & Chen WR. (2009). Optical imaging of postsynaptic odor representation in the glomerular layer of the mouse olfactory bulb. J Neurophysiol 102, 817–830.

Fukunaga I, Berning M, Kollo M, Schmaltz A & Schaefer AT. (2012). Two distinct channels of olfactory bulb output. Neuron 75, 320–329.

Gire DH & Schoppa NE. (2009). Control of on/off glomerular signaling by a local GABAergic microcircuit in the olfactory bulb. J Neurosci 29, 13454–13464.

Helassa N, Podor B, Fine A & Torok K. (2016). Design and mechanistic insight into ultrafast calcium indicators for monitoring intracellular calcium dynamics. Sci Rep 6, 38276.

Higgins NC, Escabí MA, Rosen GD, Galaburda AM & Read HL. (2008). Spectral processing deficits in belt auditory cortex following early postnatal lesions of somatosensory cortex. Neuroscience 153, 535–549.

Higgins NC, Storace DA, Escabi MA & Read HL. (2010). Specialization of binaural responses in ventral auditory cortices. J Neurosci 30, 14522–14532.

Hu XS, Ikegami K, Vihani A, Zhu KW, Zapata M, de March C, Do M, Vaidya N, Kucera G, Bock C, Jiang Y, Yohda M & Matsunami H. (2020). Concentration-dependent recruitment of mammalian odorant receptors. eNeuro.

Huang JS, Kunkhyen T, Rangel AN, Brechbill TR, Gregory JD, Winson-Bushby ED, Liu B, Avon JT, Muggleton RJ & Cheetham CEJ. (2022). Immature olfactory sensory neurons provide behaviourally relevant sensory input to the olfactory bulb. Nat Commun 13, 6194.

Igarashi KM, Ieki N, An M, Yamaguchi Y, Nagayama S, Kobayakawa K, Kobayakawa R, Tanifuji M, Sakano H, Chen WR & Mori K. (2012). Parallel mitral and tufted cell pathways route distinct odor information to different targets in the olfactory cortex. J Neurosci 32, 7970–7985.

Inagaki S, Iwata R, Iwamoto M & Imai T. (2020). Widespread Inhibition, Antagonism, and Synergy in Mouse Olfactory Sensory Neurons In Vivo. Cell Reports 31.

Kannan M, Vasan G, Huang C, Haziza S, Li JZ, Inan H, Schnitzer MJ & Pieribone VA. (2018). Fast, in vivo voltage imaging using a red fluorescent indicator. Nat Methods 15, 1108–1116.

Kapoor V, Provost AC, Agarwal P & Murthy VN. (2016). Activation of raphe nuclei triggers rapid and distinct effects on parallel olfactory bulb output channels. Nat Neurosci 19, 271–282.

Kato HK, Chu MW, Isaacson JS & Komiyama T. (2012). Dynamic sensory representations in the olfactory bulb: modulation by wakefulness and experience. Neuron 76, 962–975.

Kikuta S, Fletcher ML, Homma R, Yamasoba T & Nagayama S. (2013). Odorant response properties of individual neurons in an olfactory glomerular module. Neuron 77, 1122–1135.

Koldaeva A, Zhang C, Huang YP, Reinert JK, Mizuno S, Sugiyama F, Takahashi S, Soliman T, Matsunami H & Fukunaga I. (2021). Generation and Characterization of a Cell Type-Specific, Inducible Cre-Driver Line to Study Olfactory Processing. J Neurosci 41, 6449–6467.

Lecoq J, Tiret P & Charpak S. (2009). Peripheral adaptation codes for high odor concentration in glomeruli. J Neurosci 29, 3067–3072.

Leong LM & Storace DA. (2024). Imaging different cell populations in the mouse olfactory bulb using the genetically encoded voltage indicator ArcLight. Neurophotonics 11, 033402.

Malnic B, Hirono J, Sato T & Buck LB. (1999). Combinatorial receptor codes for odors. Cell 96, 713–723.

Martelli C & Storace DA. (2021). Stimulus Driven Functional Transformations in the Early Olfactory System. Frontiers in Cellular Neuroscience 15.

McGann JP. (2013). Presynaptic inhibition of olfactory sensory neurons: new mechanisms and potential functions. Chem Senses 38, 459–474.

Meredith M. (1986). Patterned response to odor in mammalian olfactory bulb: the influence of intensity. J Neurophysiol 56, 572–597.

Mitsui S, Igarashi KM, Mori K & Yoshihara Y. (2011). Genetic visualization of the secondary olfactory pathway in Tbx21 transgenic mice. Neural Syst Circuits 1, 5.

Nagayama S, Enerva A, Fletcher ML, Masurkar AV, Igarashi KM, Mori K & Chen WR. (2010). Differential axonal projection of mitral and tufted cells in the mouse main olfactory system. Front Neural Circuits 4.

Nagayama S, Homma R & Imamura F. (2014). Neuronal organization of olfactory bulb circuits. Front Neural Circuits 8, 98.

Nagayama S, Takahashi YK, Yoshihara Y & Mori K. (2004). Mitral and tufted cells differ in the decoding manner of odor maps in the rat olfactory bulb. J Neurophysiol 91, 2532–2540.

Niessing J & Friedrich RW. (2010). Olfactory pattern classification by discrete neuronal network states. Nature 465, 47–52.

Olsen SR, Bhandawat V & Wilson RI. (2010). Divisive normalization in olfactory population codes. Neuron 66, 287–299.

Olsen SR & Wilson RI. (2008). Lateral presynaptic inhibition mediates gain control in an olfactory circuit. Nature 452, 956–960.

Orbach HS & Cohen LB. (1983). Optical monitoring of activity from many areas of the in vitro and in vivo salamander olfactory bulb: a new method for studying functional organization in the vertebrate central nervous system. J Neurosci 3, 2251–2262.

Otazu GH, Chae H, Davis MB & Albeanu DF. (2015). Cortical Feedback Decorrelates Olfactory Bulb Output in Awake Mice. Neuron 86, 1461–1477.

Parrish-Aungst S, Shipley MT, Erdelyi F, Szabo G & Puche AC. (2007). Quantitative analysis of neuronal diversity in the mouse olfactory bulb. J Comp Neurol 501, 825–836.

Petzold GC, Hagiwara A & Murthy VN. (2009). Serotonergic modulation of odor input to the mammalian olfactory bulb. Nat Neurosci 12, 784–791.

Pinching AJ & Powell TP. (1971). The neuropil of the glomeruli of the olfactory bulb. J Cell Sci 9, 347–377.

Platisa J, Zeng H, Madisen L, Cohen LB, Pieribone VA & Storace DA. (2022). Voltage imaging in the olfactory bulb using transgenic mouse lines expressing the genetically encoded voltage indicator ArcLight. Sci Rep 12, 1875.

Polley DB, Heiser MA, Blake DT, Schreiner CE & Merzenich MM. (2004). Associative learning shapes the neural code for stimulus magnitude in primary auditory cortex. Proc Natl Acad Sci U S A 101, 16351–16356.

Polley DB, Read HL, Storace DA & Merzenich MM. (2007). Multiparametric auditory receptive field organization across five cortical fields in the albino rat. J Neurophysiol 97, 3621–3638.

Polley DB, Steinberg EE & Merzenich MM. (2006). Perceptual learning directs auditory cortical map reorganization through top-down influences. J Neurosci 26, 4970–4982.

Rabinowitz NC, Willmore BD, Schnupp JW & King AJ. (2011). Contrast gain control in auditory cortex. Neuron 70, 1178–1191.

Reisert J & Matthews HR. (1999). Adaptation of the odour-induced response in frog olfactory receptor cells. J Physiol 519 Pt 3, 801–813.

Rinberg D, Koulakov A & Gelperin A. (2006). Sparse odor coding in awake behaving mice. J Neurosci 26, 8857–8865.

Rothermel M, Brunert D, Zabawa C, Diaz-Quesada M & Wachowiak M. (2013). Transgene expression in target-defined neuron populations mediated by retrograde infection with adeno-associated viral vectors. J Neurosci 33, 15195–15206.

Rothermel M, Carey RM, Puche A, Shipley MT & Wachowiak M. (2014). Cholinergic inputs from Basal forebrain add an excitatory bias to odor coding in the olfactory bulb. J Neurosci 34, 4654–4664.

Rothermel M & Wachowiak M. (2014). Functional imaging of cortical feedback projections to the olfactory bulb. Front Neural Circuits 8, 73.

Shao Z, Puche AC, Kiyokage E, Szabo G & Shipley MT. (2009). Two GABAergic intraglomerular circuits differentially regulate tonic and phasic presynaptic inhibition of olfactory nerve terminals. J Neurophysiol 101, 1988–2001.

Shen Y, Banerjee A, Albeanu DF & Navlakha S. (2025a). An evolutionarily conserved mechanism for reformatting odor concentration in early olfactory circuits. bioRxiv, 2025.2001.2023.634259.

Shen Y, Banerjee A, Albeanu DF & Navlakha S. (2025b). An evolutionarily conserved scheme for reformatting odor concentration in early olfactory circuits. eLife Sciences Publications, Ltd.

Shusterman R, Smear MC, Koulakov AA & Rinberg D. (2011). Precise olfactory responses tile the sniff cycle. Nat Neurosci 14, 1039–1044.

Smear M, Resulaj A, Zhang J, Bozza T & Rinberg D. (2013). Multiple perceptible signals from a single olfactory glomerulus. Nat Neurosci 16, 1687–1691.

Smear M, Shusterman R, O’Connor R, Bozza T & Rinberg D. (2011). Perception of sniff phase in mouse olfaction. Nature 479, 397–400.

Stopfer M, Jayaraman V & Laurent G. (2003). Intensity versus identity coding in an olfactory system. Neuron 39, 991–1004.

Storace DA & Cohen LB. (2017). Measuring the olfactory bulb input-output transformation reveals a contribution to the perception of odorant concentration invariance. Nat Commun 8, 81.

Storace DA & Cohen LB. (2021). The mammalian olfactory bulb contributes to the adaptation of odor responses: a second perceptual computation carried out by the bulb. eNeuro.

Storace DA, Cohen LB & Choi Y. (2019). Using Genetically Encoded Voltage Indicators (GEVIs) to Study the Input-Output Transformation of the Mammalian Olfactory Bulb. Front Cell Neurosci 13, 342.

Subramanian N, Leong LM, Salemi Mokri Boukani P & Storace DA. (2025). Recent odor experience selectively modulates olfactory sensitivity across the glomerular output in the mouse olfactory bulb. Chem Senses 50.

Sutter ML & Loftus WC. (2003). Excitatory and inhibitory intensity tuning in auditory cortex: evidence for multiple inhibitory mechanisms. J Neurophysiol 90, 2629–2647.

Tian L, Hires SA, Mao T, Huber D, Chiappe ME, Chalasani SH, Petreanu L, Akerboom J, McKinney SA, Schreiter ER, Bargmann CI, Jayaraman V, Svoboda K & Looger LL. (2009). Imaging neural activity in worms, flies and mice with improved GCaMP calcium indicators. Nat Methods 6, 875–881.

Verhagen JV, Wesson DW, Netoff TI, White JA & Wachowiak M. (2007). Sniffing controls an adaptive filter of sensory input to the olfactory bulb. Nat Neurosci 10, 631–639.

Vucinic D, Cohen LB & Kosmidis EK. (2006). Interglomerular center-surround inhibition shapes odorant-evoked input to the mouse olfactory bulb in vivo. J Neurophysiol 95, 1881–1887.

Wachowiak M & Cohen LB. (2001). Representation of odorants by receptor neuron input to the mouse olfactory bulb. Neuron 32, 723–735.

Wachowiak M, Dewan A, Bozza T, O’Connell TF & Hong EJ. (2025). Recalibrating Olfactory Neuroscience to the Range of Naturally Occurring Odor Concentrations. J Neurosci 45.

Wachowiak M, Economo MN, Diaz-Quesada M, Brunert D, Wesson DW, White JA & Rothermel M. (2013). Optical dissection of odor information processing in vivo using GCaMPs expressed in specified cell types of the olfactory bulb. J Neurosci 33, 5285–5300.

Watkins PV & Barbour DL. (2008). Specialized neuronal adaptation for preserving input sensitivity. Nat Neurosci 11, 1259–1261.

Wehr M & Zador AM. (2003). Balanced inhibition underlies tuning and sharpens spike timing in auditory cortex. Nature 426, 442–446.

White EL. (1972). Synaptic organization in the olfactory glomerulus of the mouse. Brain Res 37, 69–80.

Wilson CD, Serrano GO, Koulakov AA & Rinberg D. (2017). A primacy code for odor identity. Nat Commun 8, 1477.

Xu L, Li W, Voleti V, Zou DJ, Hillman EMC & Firestein S. (2020). Widespread receptor-driven modulation in peripheral olfactory coding. Science 368.

Yu CH, Yu Y, Adsit LM, Chang JT, Barchini J, Moberly AH, Benisty H, Kim J, Young BK, Heng K, Farinella DM, Leikvoll A, Pavan R, Vistein R, Nanfito BR, Hildebrand DGC, Otero-Coronel S, Vaziri A, Goldberg JL, Ricci AJ, Fitzpatrick D, Cardin JA, Higley MJ, Smith GB, Kara P, Nielsen KJ, Smith IT & Smith SL. (2024). The Cousa objective: a long-working distance air objective for multiphoton imaging in vivo. Nat Methods 21, 132–141.

Zak JD, Reddy G, Konanur V & Murthy VN. (2024). Distinct information conveyed to the olfactory bulb by feedforward input from the nose and feedback from the cortex. Nat Commun 15, 3268.

Zak JD, Reddy G, Vergassola M & Murthy VN. (2020). Antagonistic odor interactions in olfactory sensory neurons are widespread in freely breathing mice. Nat Commun 11, 3350.

Zavitz D, Youngstrom IA, Borisyuk A & Wachowiak M. (2020). Effect of Interglomerular Inhibitory Networks on Olfactory Bulb Odor Representations. J Neurosci 40, 5954–5969.

Zhang Y, Rózsa M, Liang Y, Bushey D, Wei Z, Zheng J, Reep D, Broussard GJ, Tsang A, Tsegaye G, Narayan S, Obara CJ, Lim JX, Patel R, Zhang R, Ahrens MB, Turner GC, Wang SS, Korff WL, Schreiter ER, Svoboda K, Hasseman JP, Kolb I & Looger LL. (2023). Fast and sensitive GCaMP calcium indicators for imaging neural populations. Nature 615, 884–891.

Zhou Z, Xiong W, Zeng S, Xia A, Shepherd GM, Greer CA & Chen WR. (2006). Dendritic excitability and calcium signalling in the mitral cell distal glomerular tuft. Eur J Neurosci 24, 1623–1632.

Zhu P, Frank T & Friedrich RW. (2013). Equalization of odor representations by a network of electrically coupled inhibitory interneurons. Nat Neurosci 16, 1678–1686.

